# BMP pathway antagonism by *Grem1* regulates epithelial cell fate in intestinal regeneration

**DOI:** 10.1101/2021.01.06.425570

**Authors:** Martijn AJ Koppens, Hayley Davis, Gabriel N Valbuena, Eoghan J Mulholland, Nadia Nasreddin, Mathilde Colombe, Agne Antanaviciute, Sujata Biswas, Matthias Friedrich, Lennard Lee, Oxford IBD cohort investigators, Lai Mun Wang, Viktor H Koelzer, James E East, Alison Simmons, Douglas J Winton, Simon J Leedham

**Author notes:** These authors contributed equally to this work (Martijn AJ Koppens, Hayley Davis). **Correspondence:** Address correspondence to, Simon Leedham, Wellcome Trust Centre for Human Genetics, Roosevelt Drive, Oxford, OX3 7BN, UK.

## Abstract

In the intestine, the homeostatic effect of Bone Morphogenetic Protein (BMP) on cell fate has predominantly been inferred through pathway inactivation. Here, we assessed the impact of autocrine *Bmp4* expression on secretory cell fate. Ligand exposure reduced proliferation, expedited terminal differentiation, abrogated long-term secretory cell survival and prevented dedifferentiation. As stem cell plasticity is required for regenerative adaptive reprogramming, we spatiotemporally mapped and functionally explored BMP’s role in epithelial restitution. Following ulceration, physiological attenuation of BMP signalling arose through upregulation of the secreted antagonist, *Grem1,* from topographically distinct populations of stromal cells. Concomitant expression supported functional compensation, following *Grem1* deletion from tissue-resident fibroblasts. BMP pathway manipulation showed that antagonist-mediated BMP attenuation was obligatory, but functionally sub-maximal, as regeneration was impaired or enhanced by epithelial overexpression of *Bmp4* or *Grem1* respectively. Mechanistically, *Bmp4* abrogated regenerative stem cell reprogramming, despite a convergent impact of YAP/TAZ on cell fate in remodelled wounds.

## INTRODUCTION

The intestinal mucosa is an ideal tissue for the study of homeostatic, regenerative and pathological epithelial cell fate determination as it has stereotypical crypt-based architecture, well-defined stem cell markers and clear human disease implications. Secreted cell-signalling networks generate mucosal gradients that regulate the phenotypic response of epithelial cells migrating along the crypt^1, 2^. Bone Morphogenetic Protein (BMP) signalling is a key pathway, with a polarised gradient, maximal at the luminal surface, established through inter-compartmental crosstalk enabled by differential expression of ligands, receptors and antagonists^3, 4^. BMP ligands are predominantly secreted by the intercrypt mesenchyme and act on epithelial cells to promote differentiation. At the crypt base, exclusive expression of paracrine, ligand-sequestering BMP antagonists (e.g Gremlin1, Gremlin2, Chordin-like2) from the muscularis mucosae restricts BMP activity, and promotes epithelial stem cell activity within the crypt basal niche.

Although the physiological expression patterns of BMP pathway constituents have been mapped *in vivo*^3, 4^, the homeostatic pro-differentiation function of BMP has been predominantly inferred through pathway inactivation. Epithelial BMP receptor knockout^5^, or antagonist knock-in^6, 7^ disrupts epithelial cell fate determination and induces tumorigenesis through promotion of an aberrant stem/progenitor phenotype. The importance of BMP antagonism in promoting stem cell function was exploited in organoid system development, which require supra-physiological media concentrations of BMP antagonists for successful culture^8^. This introduces challenges for the use of organoids in assessing the nuanced physiological role of the BMP ligands.

The complexity of morphogen signalling networks permits amplification or attenuation of the biological effects of individual pathways in a dynamic and context-dependent manner. Disruption in the homeostatic morphogen balance initiates the profound multicompartmental response to injury and underpins the regenerative capacity of the intestinal epithelium. Following injury, epithelial denudation skews homeostatic epithelial-mesenchymal crosstalk and induces a localised immune response^9^. Mucosal inflammation provokes dysregulation of the fibroblastic niche, with activation of diverse stromal cell populations^10^. At a cellular level, this temporary signalling instability induces epithelial cell dedifferentiation^11^, activation of regenerative stem cells^12, 13^ and promotion of fetal epithelial cell reprogramming, partly mediated by YAP/TAZ signalling^14, 15^. The mechanisms that coordinate this profound adaptive cell reprogramming response are not well understood, and the importance of the BMP pathway is not established.

Here, we assess the impact of BMP ligand manipulation on homeostatic epithelial cell fate, spatio-temporally map the mucosal BMP signalling landscape in intestinal regeneration and assess the functional importance of BMP signalling in regulating effective wound healing.

## RESULTS

### Pan-epithelial expression of *Bmp4* reduces crypt base columnar (CBC) *Lgr5* expression

To assess the effect of BMP ligand expression on epithelial cell fate we generated a conditional *Bmp4* expressing mouse (*Rosa26*^*Bmp4*^) and crossed this with an epithelial-specific *Cre* recombinase to drive *Bmp4* ligand expression along the intestinal vertical axis (*Vil1-Cre;Rosa26*^*Bmp4*^). This abrogated the physiological gradient of BMP target gene expression in the epithelium, with *Id1*, and pSmad1,5,8, expression seen extending into the crypt bases (Fig.1a). Molecular and morphological phenotyping of the animals showed a reduction in intestinal *Lgr5* expression and CBC cell number in the epithelium^16^ (Fig.1a,b). There was subtle variation in the stromal and immune cell landscapes between mice but this did not reach statistical significance on cell quantification, except for fewer macrophages and Cd8 cells in *Vil1-Cre;Rosa26*^*Bmp4*^ small intestine (Fig.1c, Supplementary Fig.1). Notably there was no overt pathological phenotype developing in steady state mice up to 455 days after recombination, indicating homeostatic compensation which may be mediated by the enrichment of an array of BMP antagonists and signal transduction negative regulators in the *Vil1-Cre;Rosa26*^*Bmp4*^ model (Fig.1b).

**Figure 1.**
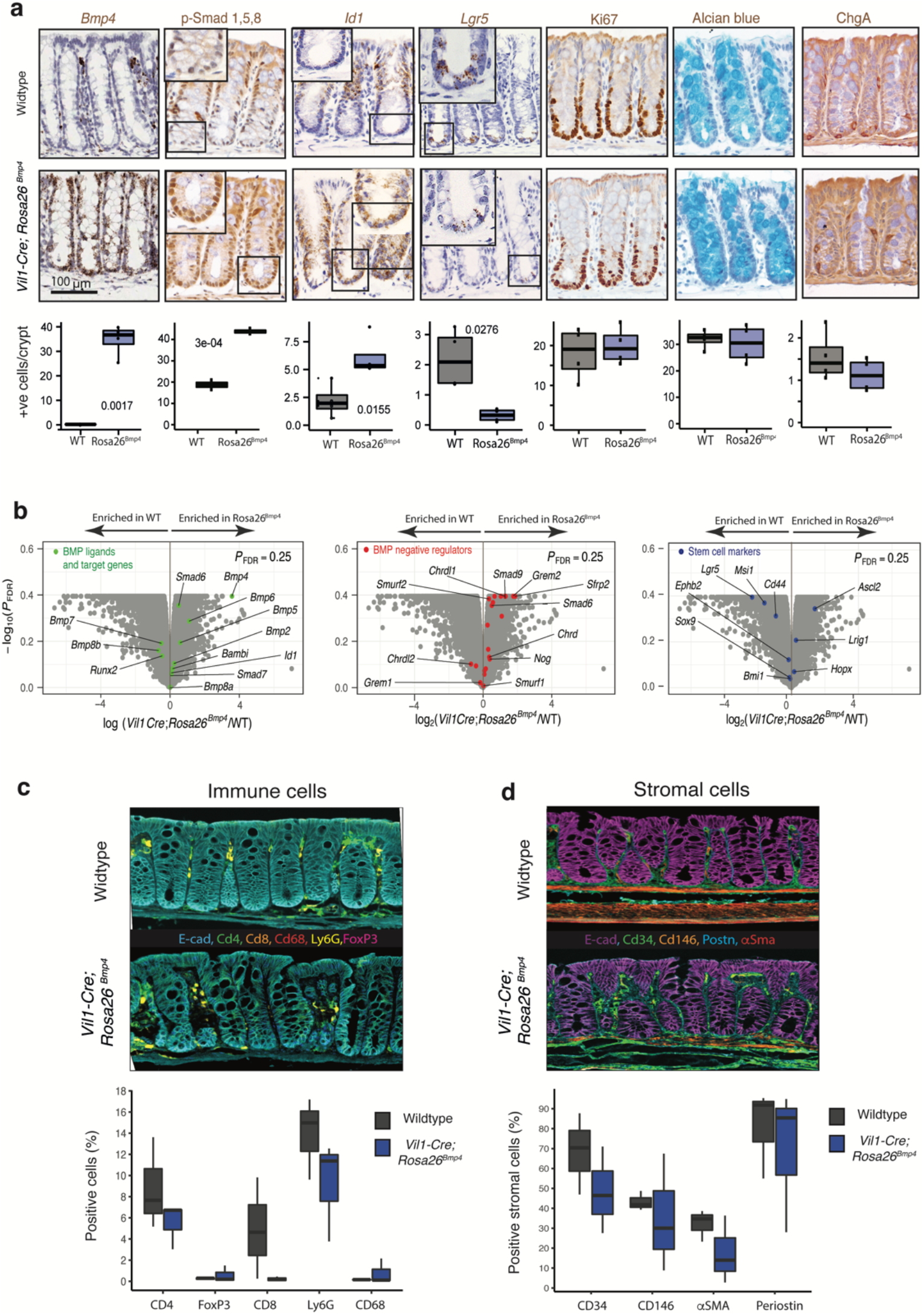
Steady-state *Vil1-Cre;Rosa26*^*Bmp4*^ mouse colonic phenotyping. **a.** ISH/IHC phenotyping and cell quantification of colon in steady state *Vil1-Cre;Rosa26*^*Bmp4*^ and control mice. (t-test, n=4 mice per genotype) **b.** Volcano plots showing differential gene expression between wildtype and *Vil1-Cre;Rosa26*^*Bmp4*^ animals. (n=3 mice per genotype) **c,d.** Multiplex IHC and cell quantification to show **c.** colonic immune and **d.** stromal cell landscapes in wildtype and *Vil1-Cre;Rosa26*^*Bmp4*^ animals (n=3 per genotype)

### Epithelial *Bmp4* ligand expression alters individual secretory cell fate determination

Beumer et al. showed that BMP ligands regulate enteroendocrine cell fate along the crypt-villus axis^17^, so we assessed the impact of autocrine expression of *Bmp4* on the fate of individual secretory progenitor cells. To do this, we used *Atoh1-CreER*^*T2*^ to induce expression of either tdTomato marker (*Atoh1-CreER*^*T2*^*;Rosa26*^*tdTom*^) or *Bmp4* (*Atoh1-CreER*^*T2*^*;Rosa26*^*Bmp4*^) in individual *Atoh1*+ve cells in steady state conditions. We then spatio-temporally tracked cells expressing either the active morphogen *Bmp4* (with ISH), or the functionally neutral tdTomato marker (with IHC), taking care to quantify and contrast spatio-temporal fate of cells within, and not directly between animal groups utilising these different methodological cell marking techniques (Fig.2a).

**Figure2.**
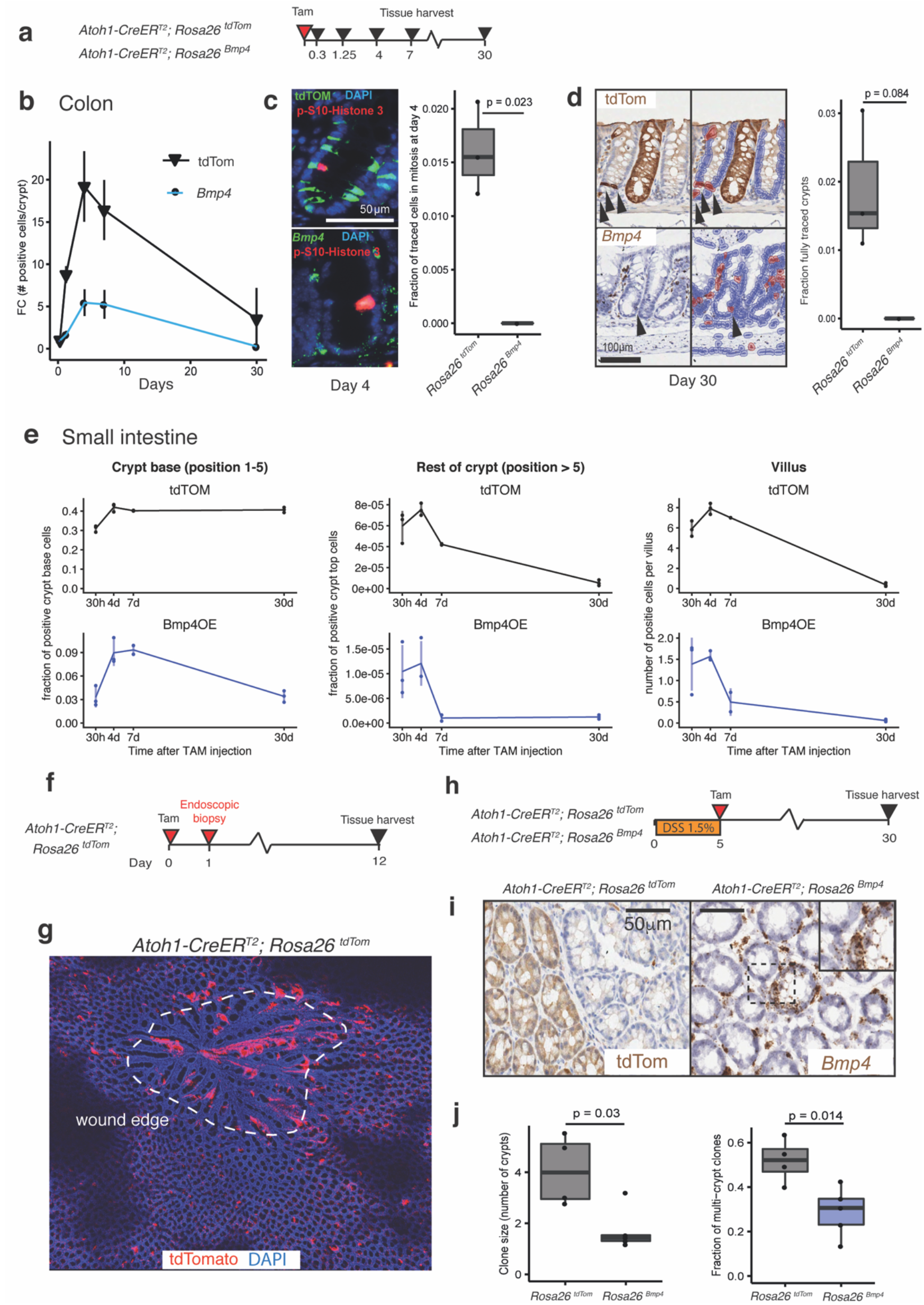
BMP ligand exposure impacts secretory progenitor cell fate. **a.** Schematic showing recombination and harvesting of homeostatic secretory cell mouse models. **b.** Number of tdTomato IHC (black lines) or *Bmp4* ISH (blue lines) stained colonic epithelial cells over time following recombination (*p* < 0.001 for Genotype, Timepoint, and Genotype: Timepoint interaction effects from a two-way ANOVA, asterisks represent statistical significance of comparisons between tdTomato and *Bmp4* at each timepoint from Tukey HSD post-hoc tests, n=3 mice per group, 7 days: n=2 mice per genotype) **c.** Co-stain of phospho-histone3 (red) with tdTomato IHC (top panel, green) or *Bmp4* ISH (bottom panel, green). Quantification of fraction of tdTomato or *Bmp4* expressing colonic cells undergoing cell proliferation at day 4 post recombination (t-test, n=3 mice per genotype). **d.** Long-lived secretory cells and lineage traced crypts 30 days after recombination, stained with tdTomato IHC or *Bmp4* ISH and automated detection of positive cells with a digital pathology platform used to exclude non-contributory stromal cell staining (QuPath). Quantification of fraction of fully traced crypts in different genotypes at day 30 (t-test, n=3 mice per genotype). *p*= 0.084 with no fully *Bmp4*-traced crypts detected. **e.** Fraction or number of tdTomato IHC or *Bmp4* ISH stained cells in the small intestinal crypt base, rest of crypt and on the villus over time following recombination (n=3 mice per group, 7 days: n=2 mice per genotype, 30 crypts and villi analysed per mouse) **f.** Schematic showing recombination and harvesting of endoscopy biopsy wounded secretory cell mouse model. **g.** *En face* section of healing endoscopic biopsy wound at day 12 (dashed white line) in *Atoh1-CreER*^*T2*^*; Rosa26* ^*tdTom*^ animals, showing streaming of recombined cells into the wound bed. **h.** Schematic showing recombination and harvesting of DSS treated secretory cell mouse models. **i.** Representative *en face* sections of colon from secretory cell models 30 days after initiation of DSS treatment, showing epithelial cell expression of tdTomato or *Bmp4* **j.** Clonal patch quantification shows an increase in multi crypt patch size and number in *Atoh1-CreER*^*T2*^*; Rosa26* ^*tdTom*^ animals following DSS treatment (t-test, n = 5 *Rosa26*^*Bmp4*^ mice, 4 *Rosa26* ^*tdTom*^ mice). Error bars represent SD. (* *p* < 0.05, ** *p* < 0.01).

In the colon, discrete secretory progenitors expressing tdTomato or *Bmp4* were identified 8 hours after recombination (Fig.2b, Supplementary Fig.2a). Cells marked with tdTomato actively proliferated (Fig.2c). The marked population peaked at day 4, with expression seen in the majority of goblet cells throughout the colonic crypts. Thereafter, the marked cell population declined, consistent with terminal differentiation, but ongoing tdTomato expression at 30 days was seen in a population of persistent colonic secretory cells, as well as a proportion of fully lineage traced crypts (Fig.2d). In *Atoh1-CreER*^*T2*^*;Rosa26*^*Bmp4*^ animals there was an absence of cell proliferation in cells expressing functionally active *Bmp4,* resulting in profound reduction in *Bmp4* marked cells at all subsequent time points (Fig.2b). At 30 days there were very few long-lived *Bmp4* expressing cells, and no fully traced crypts were seen (Fig.2d).

To assess the impact of BMP ligand on different secretory lineages, we mapped the cell position of tdTomato or *Bmp4* expressing epithelial cells in the small intestine over the same time frame (Supplementary Fig.2b). Consistent with previous findings^18^, expression of neutral tdTomato marker, was seen in crypt basal cells within 30 hours and was retained in equivalent numbers of long-lived Paneth cells for 30 days (Fig.2e, Supplementary Fig.2b). In contrast, cells at the crypt base, marked by expression of the active morphogen *Bmp4*, reduced in number after day 7, with only a third of labelled cells retained at 30 days. The proportion of *Bmp4* expressing cells in the upper crypt and villus also declined more precipitously than cells expressing tdTomato (Fig.2e; Supplementary Fig.3A). Co-staining of individual cell types showed skewed secretory cell determination at day 4 post wounding (Supplementary Fig.3b,c). These data show that in steady state, precocious exposure of secretory progenitors to BMP ligand prevents cell proliferation, expedites terminal differentiation, reduces long-lived secretory cell survival and inhibits cell dedifferentiation with resultant whole crypt lineage tracing.

### *Bmp4* expression in secretory progenitors reduces contribution to regenerative response

Dedifferentiation of *Atoh1*+ve secretory progenitor cells is critical for the regenerative response to intestinal wounding^18, 19^. To illustrate this, we undertook endoscopic biopsy wounding of recombined *Atoh1-CreER*^*T2*^*;Rosa26*^*tdTom*^ animals, collecting tissue at 11 days post wounding to assess discrete ulcer crypt dynamics (Fig.2f). Lineage traced crypts were seen surrounding the ulcer bed, with columns of tdTomato-positive epithelial cells streaming into the wound centre indicating that secretory lineage cells contributed to colonic wound healing by secondary intention^20^ (Fig.2g). To assess the impact of *Bmp4* skewing of secretory cell fate on intestinal regeneration, we quantitatively assessed crypt clonal tracing in DSS-treated A*toh1-CreER*^*T2*^*;Rosa26*^*tdTom*^ and *Atoh1-CreER*^*T2*^*;Rosa26*^*Bmp4*^ animals, using epithelial *Bmp4* or tdTomato marker expression in *en face* sections, 30 days after initiation^18^ (Fig.2h). In DSS-treated *Atoh1-CreER*^*T2*^*;Rosa26*^*Bmp4*^ animals, there was a significant reduction in multiple crypt patch size and number in contrast to the extensive clonal expansions seen in similarly treated *Atoh1-CreER*^*T2*^*;Rosa26*^*tdTom*^ mice (Fig.2j, Supplementary Fig.3d,e). This indicates that the skewing of individual secretory progenitor cell fate away from dedifferentiation reduces the contribution of *Bmp4* expressing lineages to colonic regeneration.

### Mapping the BMP signalling landscape in mouse and human colonic regeneration

Previous work in the colon and skin has suggested that physiological attenuation of the BMP pathway is required for regeneration^21, 22^ and this was consistent with our finding that autocrine ligand exposure impaired secretory cell dedifferentiation capacity. Next, we mapped the BMP signalling landscape in human inflammatory bowel disease (IBD) and two distinct mouse intestinal regeneration models (Supplementary Fig.4a,b). In human IBD, overall pathway activity, (assessed by target gene expression) was significantly decreased in IBD cases. This correlated with a corresponding increase in expression of a single intestinal secreted BMP antagonist, *GREM1* (Fig.3a). We confirmed these findings in IBD resection specimens, with upregulated *GREM1* expression seen arising from both the ulcer bed and the muscularis layers (Fig.3b). Next, we assessed pathway activity in mouse dextran sodium sulphate (DSS) colitis models and following 10Gy irradiation. The same pattern of disrupted BMP component expression was observed in murine DSS colitis, and confirmed morphologically (Fig3c,d). However, the non-ulcerating epithelial cell injury provoked by intestinal irradiation did not provoke such significant BMP pathway dysregulation (Fig.3e,f, Supplementary Fig.4c,d). Together, these data show a comparable pattern of BMP pathway suppression following barrier breach in human and mouse, with correlative stromal upregulation of *GREM1* in both human IBD and mouse DSS colitis tissue.

**Figure3.**
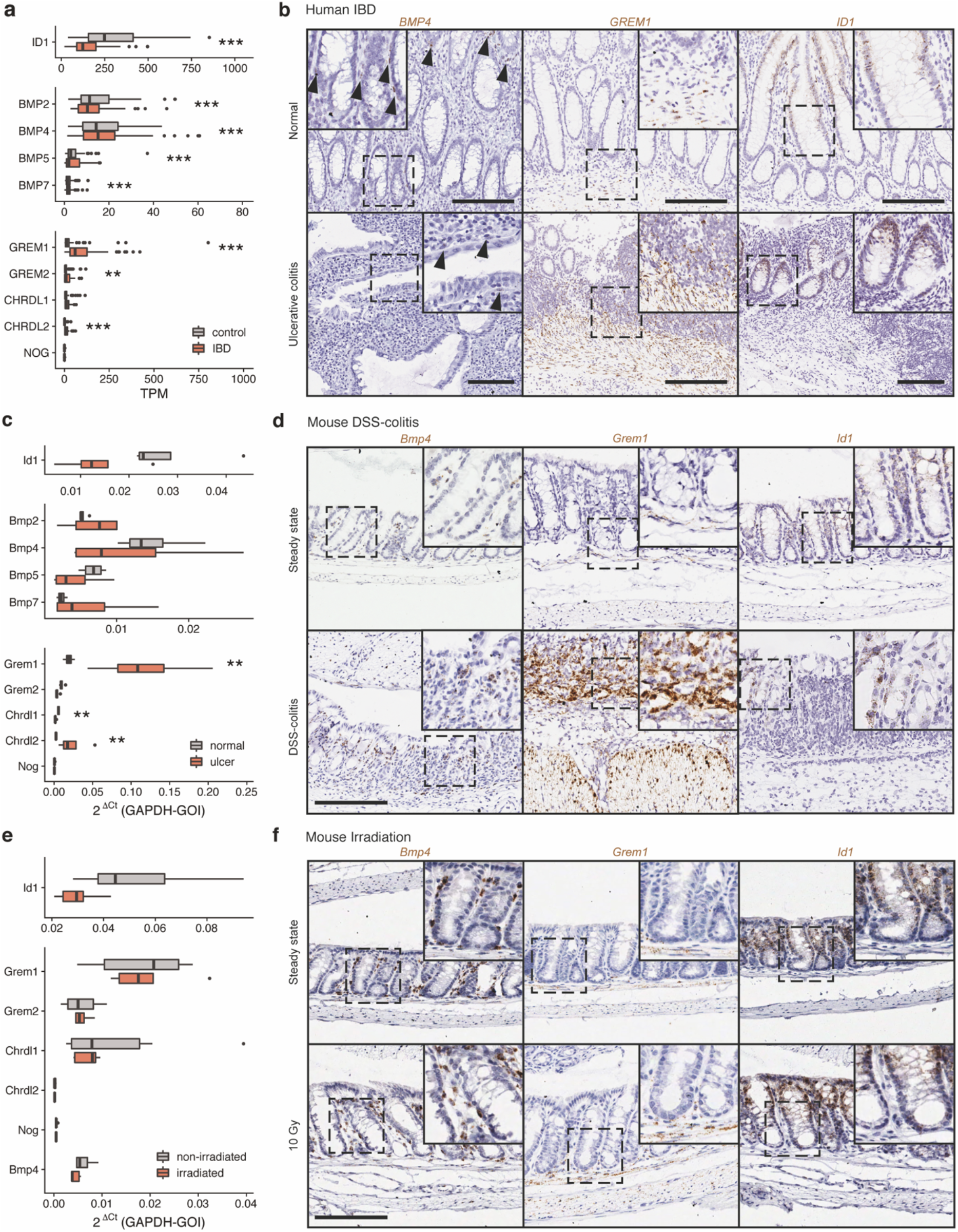
BMP signaling activity in human inflammatory bowel disease and mouse models of intestinal regeneration. **a.** Gene expression analysis of publicly-available RNAseq data from human healthy and inflammatory bowel disease tissue (GSE83687, n = 74 IBD, 60 control) shows downregulation of direct BMP target gene expression (ID1), variable impact on BMP ligand expression and corresponding strong upregulation of *GREM1* as the key intestinal BMP antagonist. **b.** *In situ* hybridization (ISH) of *BMP4, GREM1* and *ID1* in healthy human colon and severe ulcerative colitis. **c.** Colonic gene expression measured by qRT-PCR in mouse steady state and DSS colitis shows downregulation of direct BMP target gene expression (Id1), with corresponding strong upregulation of *Grem1* (*p* = 0.029 from a Mann-Whitney U test, n = 4 mice). **d.** *In situ* hybridization of *Bmp4, Grem1* and *Id1* in mouse steady state and DSS-colitis colon. **e.** Colonic gene expression measured by qRT-PCR in mouse steady state and 24 hours after 10Gy whole body irradiation, shows minimal impact of radiation damage on BMP pathway constituent expression (no compared groups were significantly different, n = 5 irradiated, 6 non-irradiated mice) **f.** *In situ* hybridization of *Bmp4, Grem1* and *Id1* in mouse steady state colon and following 10Gy irradiation. Scale bars: 200μm, magnification applies to all images, except insets. Error bars represent standard deviation (SD). Statistical differences were tested using empirical Bayes moderated two-tailed t-tests with FDR correction (a) or Mann-Whitney tests with FDR correction (c,e); (* *p* < 0.05, ** *p* < 0.01, *** *p* < 0.001).

### Topographically distinct stromal cell populations are the source of increased Grem1 expression

Having identified profound upregulation of a single BMP antagonist in mouse DSS colitis models, we used a dual morpho-molecular approach to characterise the *Grem1* expressing cell population(s). First, we excluded *Grem1* expression arising from immune and vascular cells (Fig.4a, Supplementary Fig.5a,b). We then used dual ISH/IHC to spatially segregate *Grem1* expressing mesenchymal cells into two topographically distinct groups - αSMA-marked muscularis mucosae/propria cells, and a previously demonstrated, inflammation-expanded, heterogenous population of stromal cells in the ulcer bed, broadly marked by Podoplanin (PDPN/gp38)^23,24^ (hereafter referred to as wound-associated stromal cells, WASC) (Fig.4a). We used published stromal scRNA from DSS colitis models^10^, in combination with dual ISH/IHC, to assess the co-expression and spatial distribution of established and functionally relevant, stromal sub-populations within these broader mesenchymal groups (Fig.4b, Supplementary Fig.5). We were able to find overlap of *Grem1* expression with some key morphogens and cytokines implicated in colonic regeneration, with stromal subsets co-expressing *Grem1* and *Rspo3*^25^, *Ptgs2* (COX2)^26^ and *Il33*^27^. However, there was no significant overlapping expression with other stromal cells previously identified as subset markers^10^ or mesenchymal Wnt ligand sources including *Foxl1*^28^, *Gli1*^29^ and *Wnt5a*^20^ (Supplementary Fig.5c-e).

**Figure4.**
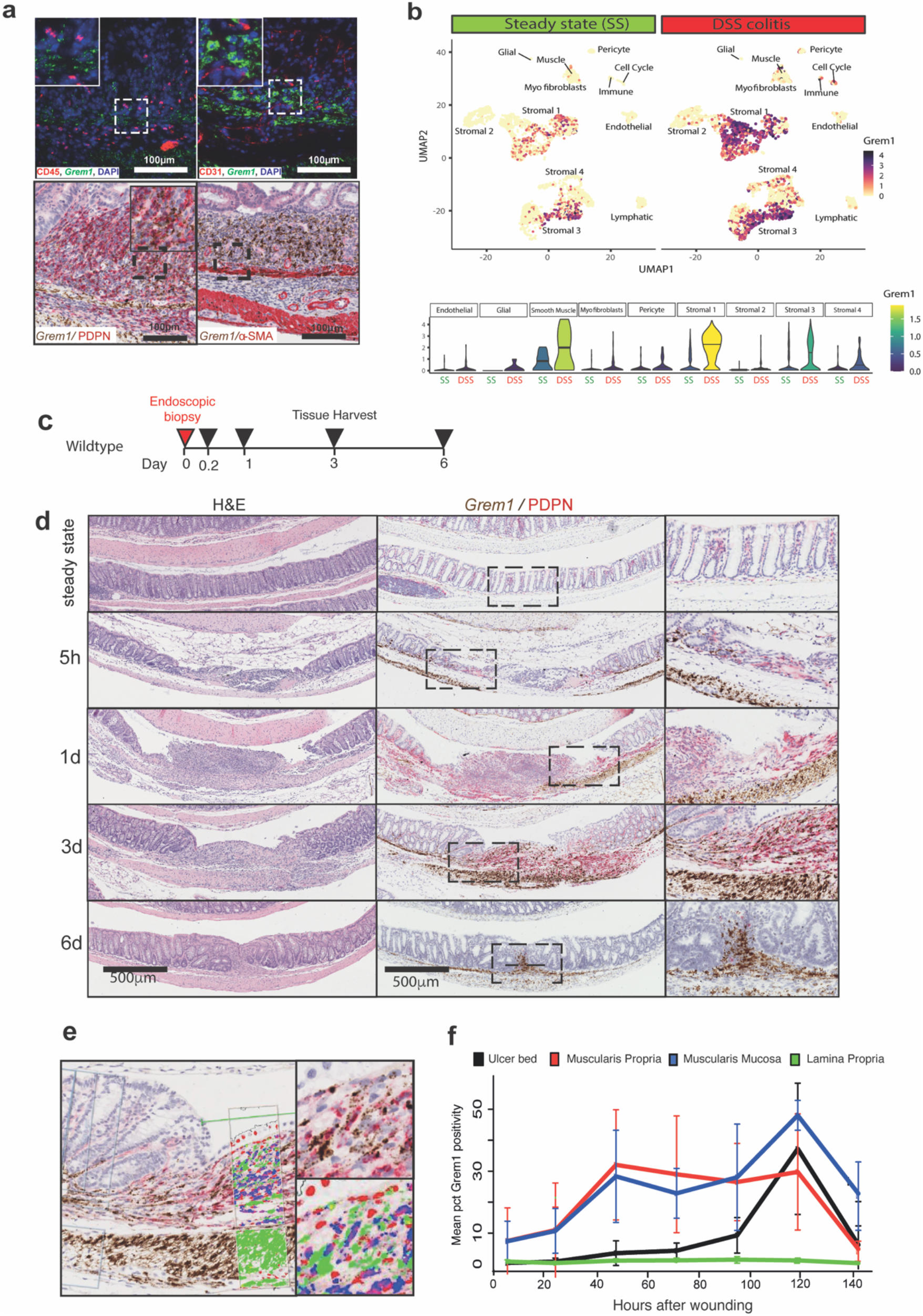
Two stromal cell populations upregulate *Grem1* in response to colonic injury. **a.** *in situ* hybridisation for *Grem1* expression with concomitant immunohistochemical co-staining for stromal cell identification. There was no overlap of fluorescent expression of *Grem1* mRNA (green) with leucocytes marked by CD45 (red) or endothelial cells marked by CD31 (red). Overlapping chromogenic ISH for *Grem1* (brown) was seen with both ulcer bed stromal cells marked with podoplanin (PDPN) IHC (red), and muscularis mucosae/propria cells marked by αSMA IHC stain (red) **b.** scRNA U-MAP plots showing upregulation and diversification of *Grem1* expressing stromal cell populations in mouse steady state (SS) and DSS colitis. Violin plots show *Grem1* expression in steady state (SS) and DSS colitis in the mesenchymal cell subsets identified by Kinchen et al.^10^ **c.** Schematic of biopsy schedule and tissue harvesting for endoscopic biopsy wound mice. **d.** H&E and *Grem1* chromogenic ISH (brown) co-stained with PDPN IHC (red) of biopsy-wounded WT mice at 5 hours, 1 day, 3 days and 6 days after wounding. Scale bar: 500μm. **e.** Digital pathology false colour markup of a *Grem1*/PDPN co-staining image at 3 days after wounding. **f.** Spatio-temporal quantification of *Grem1*-positivity in different colon tissue layers in and within 0.5mm of wound. Error bars represent SD.

DSS colitis is a simple and reproducible model for provoking a colonic regenerative response, but is not suitable for temporal interrogation, as ulceration initiation can occur at an unknown point during the 5-7 day administration period. To assess spatio-temporal dynamics of *Grem1* expression from stromal cells following injury, we undertook endoscopy-guided colon biopsy wounding (Fig.4c). Strikingly, *Grem1* expression was upregulated in the muscularis mucosa/propria of the ulcer environs as early as 5 hours after injury and persisted to at least day 6 post wounding (Fig.4d). Podoplanin cell staining was seen in the ulcer bed within 24 hours, however very little *Grem1* expression was observed from WASC until >48 hours after injury induction. These results indicated that increased expression of *Grem1* from the muscularis and WASC populations was both spatially and temporally distinct, but had a cumulative effect of generating a rapidly induced, and sustained increase in *Grem1* expression in the vicinity of the wound (Fig.4e,f).

### Genetic manipulation of BMP signalling activity affects colonic regenerative capacity

To assess the importance of secreted BMP antagonism in suppressing physiological signalling in colitis, we used *Cagg-CreER*^*T2*^*;Grem1*^*fl/fl*^ mice to knockout intestinal stromal *Grem1* expression^7, 30^ (Fig.5a). Although efficient recombination and knockout was seen in the αSMA-positive muscularis cells throughout the colon, WASC continued to robustly express *Grem1* in colonic ulcer beds (Fig.5b). Consequently, any potential detrimental effect of the conditional Grem1 knockout on wound healing fell below our significance threshold. (Fig.5c,d,f). Multiplex IHC showed no significant quantifiable variability in stromal cell marker expression in this mouse genotype (Fig.5d,e), so these data indicate that WASC arise from activation of stromal cell populations that are not effectively recombined in *Cagg-CreER*^*T2*^ animals^31^. Concomitant expression from these populations can partially compensate for *Grem1* knockout in tissue-resident αSMA positive muscularis cells.

**Figure5.**
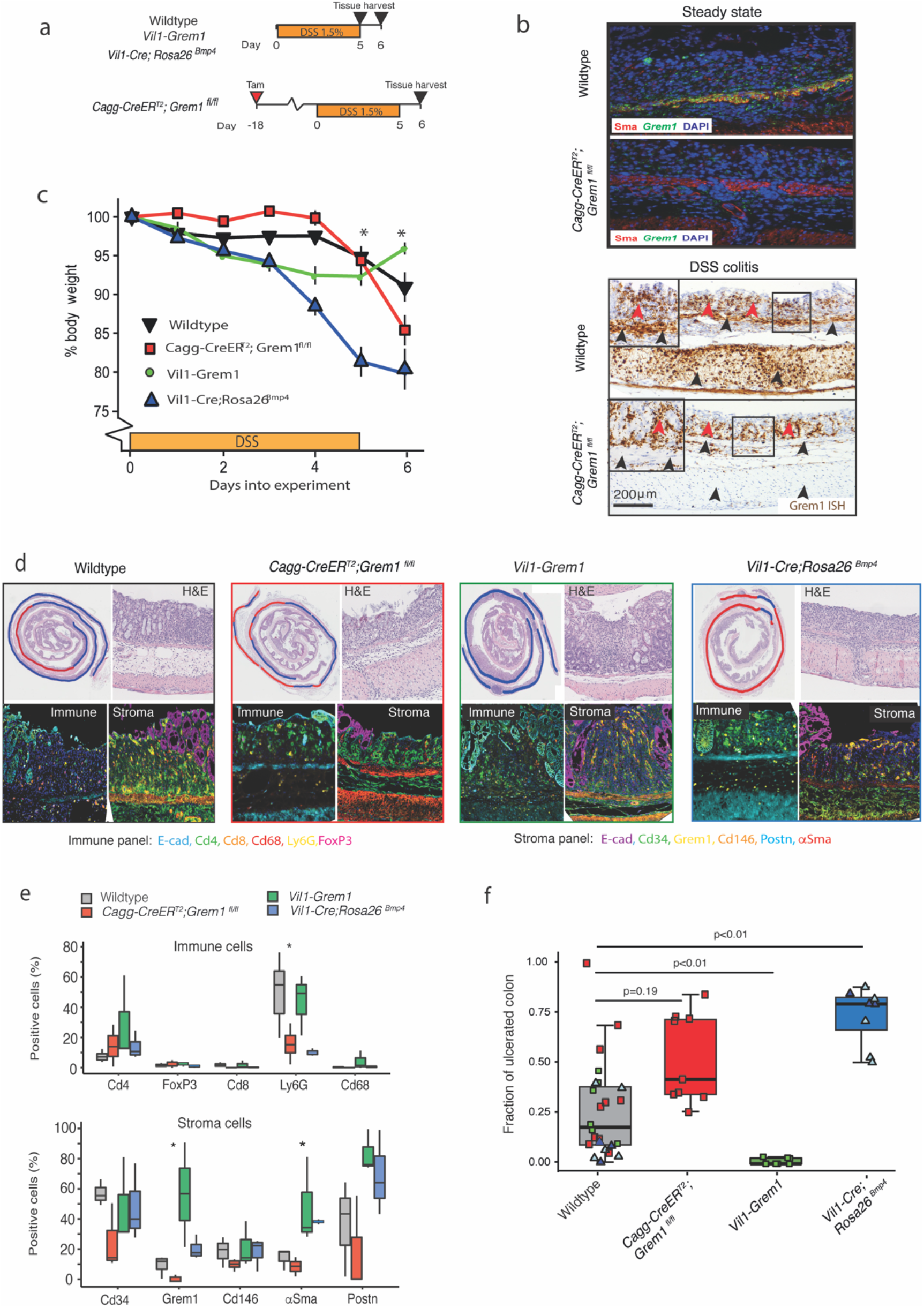
Functional impact of BMP manipulation on tissue regeneration. **a.** Schematic of recombination (where applicable), DSS administration and tissue harvesting schedule for constitutive and inducible mouse models. **b.** In steady state, *Grem1* ISH (green) and αSMA IHC (red) showing clear co-localisation of expression in the muscularis mucosa of wildtype animals and efficient knockout of *Grem1* expression in *Cagg-CreER*^*T2*^*;Grem1*^*fl/f*^^l^ animals. In DSS colitis, successful knockout of *Grem1* expression is seen from the tissue-resident muscularis mucosa and propria (black arrowheads), but continued *Grem1* expression (brown stain) from the wound-associated stromal cells persists (red arrowheads). **c.** Body weight of WT (black line), *Cagg-CreER*^*T2*^*;Grem1*^*fl/f*^^l^ (red line), *Vil1-Grem1* (green line) and *Vil1-Cre;Rosa26*^*Bmp4*^ (blue line) mice over time as percentage of body weight at start of DSS-treatment (asterisks show statistical significance from t-tests comparing WT and *Vil1-Cre;Rosa26*^*Bmp4*^ at each timepoint, n=6-8 mice per genotype; 5/8 Rosa26-Bmp4 mice were killed at day 5 along with 5/8 WT littermates). Error bars represent S.E.M. **d.** Representative H&E images of DSS-induced ulceration with blue and red curves demarcating the extent of normal and denuded colon respectively. Multiplex IHC stain to show immune and stromal cell landscapes of DSS induced ulcers in wildtype (grey box), *Cagg-CreER*^*T2*^*;Grem1*^*fl/f*^^l^ (red box), *Vil1-Grem1* (green box) and *Vil1-Cre;Rosa26*^*Bmp4*^ mice (blue box) **e.** Cell quantification of multiplex IHC immune and stromal cell stain in DSS ulcers between genotypes (* p<0.05, ANOVA, n=3 per genotype). **f.** Fraction of ulcerated colon in different animal genotypes. Control coloured dots represent control littermates for the individual genotype experiments. Light blue triangles are 5 (out of 8) *Vil1-Cre;Rosa26*^*Bmp4*^ animals that exceeded weight loss limits and had to be killed at 5 days; an equal number of wildtype littermates were killed as time point controls. Dark blue triangles are wildtype or *Vil1-Cre;Rosa26*^*Bmp4*^ animals that reached the 6 day experimental endpoint (3/8). Error bars represent SD. (* *p* < 0.05, ** *P* < 0.01, t-test, n=6-9 mice per genotype)

This compensation arising from the heterogeneity of *Grem1* expressing mesenchymal cells, meant that to assess the functional effects and therapeutic potential of BMP pathway manipulation in regeneration, we needed to maximise agonism and antagonism of the pathway through epithelial overexpression of *Bmp4* (*Vil1-Cre;Rosa26^*Bmp4*^)* or *Grem1* (*Vil1-Grem1)*^7^ respectively (Fig.5a). Although *Vil1-Cre;Rosa26*^*Bmp4*^ mice had no steady-state pathological phenotype, these animals lost a significant amount of weight following DSS treatment, requiring early sacrifice in some cases (Fig.5c). Mice exhibited a dramatic phenotype, with complete distal colonic epithelial loss (Fig.5d,f). In contrast, DSS treatment of *Vil1-Grem1* animals had minimal colitogenic impact, with barely detectable macroscopic ulceration (Fig.5d,f), although the colonic epithelium was hyperplastic^7^. Together these data demonstrate that manipulation of the physiological mucosal BMP gradient, through epithelial ligand or antagonist expression significantly alters intestinal regenerative capacity. An excess of BMP ligand profoundly impairs epithelial restitution, whilst maximising pathway antagonism through ectopic *Grem1* expression, enhances wound healing at the expense of epithelial hyperplasia.

### BMP antagonism and YAP/TAZ signalling convergently impact epithelial cell fate

To assess the impact of manipulated BMP signalling on epithelial spatio-temporal dynamics in wound healing we turned back to the endoscopic biopsy model (Supplementary Fig.6a). Biopsy ulcers in *Vil1-Grem1* animals were rapidly covered by wound-associated epithelium (WAE) (Fig.6a), whereas epithelial *Bmp4* expression in *Vil1-Cre;Rosa26*^*Bmp4*^ mice delayed coverage by WAE and retarded closure (Fig.6b, Supplementary Fig.6b). Next, we assessed stem and proliferating cell dynamics. Loss of CBC cells has been reported following intestinal injury^32^ and *Lgr5*-expressing cell depletion was seen in ulcer-adjacent crypts immediately after wounding. CBC recovery from day 3 was enhanced by epithelial *Grem1* expression in the *Vil1-Grem1* model and delayed in *Vil1-Cre;Rosa26*^*Bmp4*^ (Fig.6c, Supplementary Fig.6c). Following an appropriate wound-induced spike in cell division in ulcer-adjacent crypts at 24 hours in all genotypes, proliferation slowed in *Vil1-Cre;Rosa26*^*Bmp4*^ animals (Fig.6d, Supplementary Fig.6d), consistent with delayed reconstitution of CBC populations. We confirmed enrichment of fetal signatures^15^ in endoscopic biopsy wounds (Fig.6e) and showed that epithelial *Bmp4* ligand prevented expression of this reprogramming signature in 3-day old wounds (Fig.6f). Ayyaz *et al.* showed that activation of *Lgr5*-negative regenerative stem cells was marked by expression of the molecular chaperone Clusterin *(Clu)*^12^, a gene that is also enriched in the fetal signature. We used multicolour ISH to track temporal expression of this cell marker over time following biopsy. In WT and *Vil1-Grem1* animals, *Clu* expression was rapidly activated in WAE and ulcer-adjacent crypts, and the *Clu*+ve cell population expanded over time. (Fig.6g,h, Supplementary Fig.6e). However, *Vil1-Cre;Rosa26*^*Bmp4*^ animals exhibited significant delay in *Clu* expression which did not appear until >day 3 following injury. Together these data show that post-wounding regenerative stem cell activation, marked by *Clu* expression and characterised by fetal gene signatures, was enhanced by BMP antagonism and temporally retarded by autocrine epithelial *Bmp4* signalling.

**Figure6.**
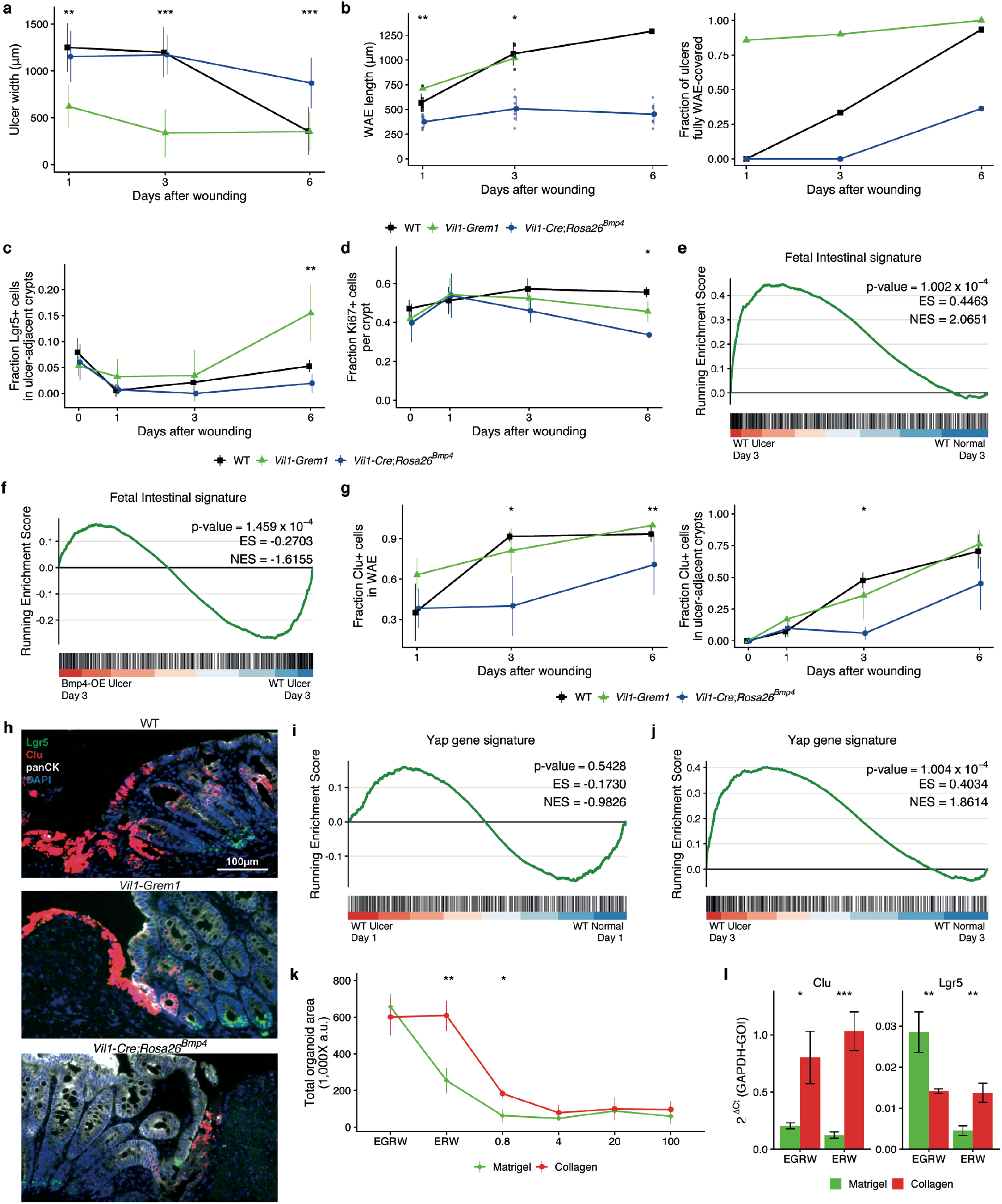
BMP manipulation affects epithelial adaptive response in wound healing. Microscopic appearance of ulcers in different genotypes over time following endoscopic wounding - **a.** Ulcer width, **b.** wound-associated epithelium (WAE) length and the fraction of ulcer covered by WAE (n =6-15 ulcers). Wounds completely covered with WAE were excluded from WAE length measurements. **c.** fraction of *Lgr5* positive cells (n = 3-6 mice per group), and **d.** Ki67 positive proliferating cells in ulcer-adjacent crypts in different genotypes over time (n = 3-5 mice per group). **e.** Gene set enrichment analysis (GSEA) showing enrichment of fetal intestinal signature in the endoscopy biopsy wound milieu at day 3 and **f.** negative enrichment of fetal intestinal signature in the biopsy wounds of *Vil1-Cre;Rosa26*^*Bmp4*^ mice in comparison with wounded wildtype animals at day 3. **g.** Regenerative stem cells, marked by clusterin (*Clu*) staining in **i.** WAE and **ii.** ulcer-adjacent crypts in different genotypes, over time (n = 3-6 mice per group). In all line plots, line colours represent: wildtype (black), *Vil1-grem1* (green) and *Vil1-Cre; Rosa26*^*Bmp4*^ (blue). **h.** Representative images of ISH for *Lgr5* (green), *Clusterin* (red) with co-stain IHC for pan-cytokeratin (white) and DAPI (blue) in 3 day old endoscopy wounds in different genotype animals. Scale bar: 100μm. **i.** Gene set enrichment analysis showing no enrichment of YAP signature in acute biopsy wounds in wildtype animals at day 1, but **j.** significant enrichment of YAP signature in biopsy wound ulcers by day 3 as the wound bed remodels. **k.** Organoid survival when grown in matrigel (green line) or collagen (red line) with media containing variable recombinant proteins and increasing doses of recombinant BMP4 (n = 3 experiments, n=2 for 20ng/mL). **l.** qRT-PCR of crypt base columnar (*Lgr5*) and regenerative stem cell gene expression (*Clu*) in organoids grown in matrigel or collagen (n = 3 experiments) and with variable media constituents (E: Epidermal growth factor, G: GREM1, R: RSPO1, W: WNT3A). Error bars represent SD. Statistical differences were tested using a Kruskal-Wallis test (a-d, f, h) or a Student’s t-test (I); (* *P* < 0.05, ** *P* < 0.01, *** *P* < 0.001)

We noted that delayed, but not entirely eliminated activation of *Clu*-positive regenerative stem cells following day 3 in *Vil1-Cre;Rosa26*^*Bmp4*^ animal wounds, occurred despite ongoing *Bmp4* expression. YAP/TAZ signalling plays a role in regulating adaptive reprogramming following barrier breach^14, 15^. We were able to confirm enrichment of YAP/TAZ signatures from mature endoscopic wounds at day 3 (Fig.6j), but saw no significant upregulation in biopsy wounds at day 1 (Fig.6i). Given this temporal lag in YAP/TAZ activation, we postulated that matrix remodelling in mature wounds might independently contribute to delayed activation of regenerative (*Clu*+ve) stem cells in *Vil1-Cre;Rosa26*^*Bmp4*^ animals. To assess this we turned to organoid models, as it has been shown that growth in collagen matrix simulates a remodelled wound bed, by activating epithelial YAP/TAZ signalling, and enriching for fetal signature expression^15^. We cultured mouse intestinal organoids in both matrigel and collagen and assessed the media BMP antagonist requirement (GREM1), and the tolerance of media BMP4 ligand in both conditions. Organoids grown in Matrigel were enriched for *Lgr5* expression, had an obligatory culture requirement for media BMP antagonist (GREM1) supplementation, and growth was inhibited by very low concentrations of BMP4 ligand (Fig.6k,l, Supplementary Fig.6f). In contrast, organoids grown in collagen were enriched for *Clu* over *Lgr5*, had acquired BMP antagonist independence, as they no longer required media GREM1 inclusion, and were less sensitive to growth suppression by low doses of BMP4 ligand. Thus, organoid growth in collagen-enriched matrix can elicit a regenerative stem cell response through YAP/TAZ activation, and this abrogates the absolute requirement for BMP antagonism in organoid culture.

Altogether these results show that BMP manipulation *in vivo* can skew the dynamic temporal response to an acute wound, with exaggerated antagonism in the *Vil1-Grem1* model enhancing regenerative stem cell activation, CBC reconstitution, cell proliferation and epithelial restitution. In contrast, epithelial *Bmp4* ligand overexpression delays and abrogates the epithelial reprogramming response, with impact on regenerative capacity. Although excessive BMP ligand delays the epithelial response, it does not entirely abrogate regenerative stem cell activation in remodelled wounds, and we show a temporally staggered but functionally convergent contribution from YAP/TAZ signalling to adapt epithelial cell fate across the duration of wound healing.

## DISCUSSION

Current IBD therapeutics reduce provoking inflammation^33^ with limited focus on enhancing the mucosal regenerative capacity, as the mechanisms that regulate epithelial restitution are incompletely understood. Here, we have used a combination of models to map and functionally assess the impact of BMP pathway manipulation on epithelial cell fate in homeostasis, and in the regenerative response to wounding. In steady state, autocrine epithelial *Bmp4* expression skews secretory cell fate, expediting terminal differentiation and inhibiting progenitor cell dedifferentiation. As regeneration is underpinned by rapid epithelial adaptive cell reprogramming there is a physiological requirement for attenuation of BMP signalling following injury. We show that this attenuation is mediated through upregulated expression of the secreted antagonist *Grem1,* from topographically distinct stromal cell populations. Cumulative *Grem1* expression from independent mesenchymal cell populations contributes to rapidly initiated yet sustained BMP antagonism localised to the injury environs, and permits capacity for functional compensation. Despite profound physiological stromal *Grem1* upregulation, we demonstrate functionally sub-maximal BMP antagonism following colonic ulceration, as ectopic epithelial *Grem1* expression in the *Vil1-Grem1* model markedly expedited colonic epithelial regeneration, highlighting the potential for BMP pathway manipulation in future IBD therapeutics. However, it is important to note that the enhancement of wound healing capacity through permanent epithelial *Grem1* exposure, comes at the longer term expense of generating a hyperplastic, and ultimately pro-tumorigenic epithelial environment^7^.

The multicompartmental mucosal response to barrier breach is complex. Different studies have highlighted the involvement of other mechanisms in regulating intestinal regeneration, such as metabolic stress^13^, mechanotransduction^14, 15^ and numerous other secreted cell signalling pathways, such as *TGFβ*^20^ and Wnt signalling^20, 25, 34^. Here, we show variable requirement for BMP suppression in regulating gut regeneration following different modes of injury, with more pronounced disruption of mucosal BMP signalling provoked by ulceration than seen following intestinal irradiation. This reflects differences between the injurious stimuli, with irradiation predominantly damaging the epithelial cell compartment, whilst barrier breach invokes a multi-compartmental tissue response. The use of spatio-temporal mapping to assess variability in cell signalling disruption following induction of intestinal injuries can help to triangulate the role of different pathways. Using this technique, alongside organoid models, we demonstrate a temporally-staggered, but functionally convergent effect of paracrine BMP antagonism and YAP/TAZ activation on promotion of the epithelial adaptive cell response in the ulcer milieu. Given the complexity of the multicompartmental response to wounding it is likely that multiple different, but frequently intersecting pathways act in a convergent fashion to alter epithelial cell fate determination. Understanding this synergism, and any functional redundancy in the system will be vital for development of drugs that therapeutically manipulate the intestinal regenerative response.

The intestinal mucosa has evolved to generate an effective and rapid response to injury, through temporary relaxation of stringent homeostatic cell fate control. Epithelial denudation, stromal activation and immune cell infiltration induces transient disruption of physiological, polarised signalling gradients which drives surrounding epithelial cell dedifferentiation and adaptive reprogramming. As wounds heal, barrier restitution, generation of new crypt architecture and dissolution of immune and activated stromal cell infiltrate allows re-imposition of graduated cell signalling and re-enforces regimented cell fate determination along the crypt axis. Here we show that excessive BMP ligand exposure impacts epithelial cell dedifferentiation capacity, hence intestinal regeneration is dependent on physiological attenuation of this homeostatic signal, which occurs through time-limited upregulation of the secreted antagonist, GREM1. As we have previously shown, permanent over-expression of the same antagonist is pro-tumorigenic in humans^35^ and mice^7^, and this illustrates the fine line between dynamic, temporary disruption of signalling networks to physiologically adapt epithelial cell fate in regeneration and the pathological co-option and corruption of the same pathways in neoplasia.

## METHODS

### Animal Models

All animal experiments were carried out in accord with the guidelines of the UK Home Office, under the authority of a Home Office project license approved by the Animal Welfare and Ethical Review Body at the Wellcome Centre for Human Genetics, University of Oxford. All mice were housed in a specific-pathogen-free (SPF) facility, with unrestricted access to food and water, were not involved in any previous procedures. All strains used in this study were maintained on C57BL/6J background for ≥6 generations. All procedures were carried out on mice of at least 6 weeks of age, both male and female.

### Generation of *Rosa26*^*Bmp4*^ *mice*

To generate *Rosa26*^*Bmp4*^ *mice, a* Bmp4 cDNA cassette was cloned into the integrase mediated cassette exchange vector (CB93) and transfected into RS-PhiC31 ES cells. Recombinant clones were obtained which harbour the Bmp4 cDNA transgene positioned within the Rosa26 locus, allowing for Cre-dependent activation of transgene expression. Recombinant clones were injected into blastocysts and chimeras generated (trangenics core, Wellcome Centre for Human Genetics). Chimeras were crossed with wild-type C57BL/6J mice to obtain F1 heterozygotes. The following primers were used for genotyping (F rbGpaF1 CAGCCCCCTGCTGTCCATTCCTTA, R Rosa3HR CGGGAGAAATGGATATGAAGTACTGGGC) and (F Caggs CAGCCATTGCTTTTATGGT, R Ex Neo2 GTTGTGCCCAGTCATAGCCGAATAG).

### Treatment of Animals

Induction of CreER^T2^ in animals was carried out using the free base tamoxifen (Sigma) dissolved in ethanol/oil (1:9). To obtain *Grem1* knockout mice, 6 weeks old *Cagg-CreER*^*T2*^*;Grem1*^*fl/fl*^ mice were injected with tamoxifen intraperitoneally, 1 mg daily, for 5 days. For lineage tracing following biopsy wounding (*Atoh1CreER*^*T2*^*;Rosa26*^*tdTomato*^) recombination was induced by a single dose of 3 mg tamoxifen, biopsy-wounding was performed 24 hours post tamoxifen injection. Biopsy wounding procedure was performed on isofluorane-anesthetized mice. First lubricant was applied to its anus, then a miniature rigid endoscope (1.9mm outer diameter, Karl Storz) was inserted 2-3cm into the rectum. Three biopsies were taken from well-separated areas in the colon wall using forceps (1mm, 3 French). The mice were killed at various time points post wounding. SI injury was induced by exposing animals to whole-body irradiation (10 Gy) performed using an IBL-637 irradiator with a Caesium-137 source. Irradiation was delivered over 652 seconds at 0.92 Gy/Min. For DSS-induced colitis, mice were treated with drinking water containing 1.5% DSS (MP Biomedicals) for a period of 5 days, after which their water was changed back to normal water to recover for one day, after which they were sacrificed. For experiments using lineage tracing in DSS-wounded mice, a single dose of 3 mg tamoxifen via intra-peritoneal injection was given immediately after 5d 1.5% DSS treatment in the drinking water. The mice were weighed daily and checked for signs of discomfort (e.g. hunching, pale feet). The mice were killed when their weight dropped below 80% of their weight at the start of the DSS treatment, or when they showed other clear symptoms of discomfort.

### Human subjects

Human pathological samples were used from normal tissue distant to tumour resections, or from ulcerative colitis specimens, taken from bowel resections for acute severe colitis following ethical approval and individual informed consent (MREC 16/YH/0247).

### RNA extraction and gene expression analysis

For gene expression analysis by qPCR, tissues and cells were lysed and RNA was isolated using the RNeasy Mini kit (QIAGEN). Organoids were first extracted from collagen matrix by incubating with collagenase VIII (Sigma-Aldrich, 0.6ug/uL in medium) for 5 minutes, or from Matrigel by washing with ice-cold medium, before lysis. 1cm of distal colon (irradiation experiment) was homogenised using a handheld homogeniser (Polytron, Kinematica). RNA was DnaseI treated (ThermoFisher Scientific). cDNA was generated using the High Capacity cDNA Reverse Transcription Kit (ThermoFisher Scientific). To roughly normalize samples within an experiment, the same amount of RNA was taken per sample to make cDNA. Quantitative RT-PCR (qRT-PCR) was performed on the Applied Biosystems QuantStudio 6 Flex Real-Time PCR System using Fast Universal PCR Master Mix and TaqMan Gene Expression assays (Applied Biosystems), listed in Table S1.

### RNA-sequencing

Colon ulcers and distant normal colon tissue were excised from biopsy-wounded wild-type and *Vil1-Cre;Rosa26*^*Bmp4*^ mice at 1 day and 3 days post-wounding and stored in RNAlater (ThermoFisher) at −80°C prior to extraction. Total RNA was extracted using the RNeasy Mini Kit (Qiagen), with on-column DNAse treatment. RNA concentrations and purity were assessed using a NanoDrop One (ThermoFisher). Samples with A260/A280 ratio < 2 were subjected to further DNAse treatment using DNAse I, Amplification Grade (18068015, ThermoFisher). RNA concentrations were measured using the Qubit RNA HS Assay Kit (Q32855, ThermoFisher). RNA quality was assessed on a 2100 Bioanalyser Instrument (Agilent). Library construction and sequencing were performed at the Oxford Genomics Centre. Total RNA quantity and integrity were assessed using the Quant-IT RiboGreen RNA Assay Kit (Invitrogen, Carlsbad, CA, USA) and the Agilent Tapestation. mRNA purification, double-stranded cDNA generation and library construction were performed using NEBNext Poly(A) mRNA Magnetic Isolation Module (E7490, New England Biolabs) and NEBNext Ultra II Directional RNA Library Prep Kit for Illumina (E7760L, New England Biolabs) with in-house adapters and barcode tags (using dual indexing, based on doi:10.1186/1472-6750-13-104). The concentrations used to generate the multiplex pool were determined by a Quant-iT PicoGreen dsDNA Assay (P7589, Invitrogen). The final size distribution of the pool was determined using a Tapestation system (Agilent), and quantified using a Qubit assay (Thermofisher). Libraries were sequenced on the NovaSeq 6000 (Illumina) as 150-bp paired-end reads.

### Data and Code Availability

The mouse RNAseq data generated for this study have been deposited in the European Nucleotide Archive (ENA) at EMBL-EBI under accession number PRJEB40623.

### Bioinformatic analysis of RNA sequencing

RNA sequencing data from human inflamed colon tissue from IBD patients and non-inflamed colon tissue from non-IBD controls (Peters, et al., 2017) were downloaded from the European Nucleotide Archive (accession number PRJNA326727). Raw sequence reads were subjected to adapter trimming using *BBduk* (*BBTools* ver. 38.46). Trimmed reads were aligned to the GRCh38 build of the human reference (for human colon tissue data) or to the GRCm38 build of the mouse reference (for mouse ulcer data) using *STAR* (ver. 2.7.0f). Ensembl 96 annotations were used for alignment and subsequent quantifications. Gene expression was quantified using *RSEM* (ver. 1.3.1). The processed RNAseq data was analysed in the R statistical environment (ver. 3.6.1). Differential expression analyses were performed using the *limma* package (ver. 3.40.6). Gene Set Enrichment Analyses were performed using the *fgsea* package (ver. 1.10.1). The YAP/TAZ gene signature includes genes that had > 2-fold upregulation between wild-type and YAP-overexpressing intestinal epithelial cells ^14^, while the Fetal Intestinal gene signature includes genes that were significantly upregulated > 2-fold (FDR < 0.05) in fetal intestinal epithelial cell cultures over adult intestinal epithelial cell cultures^15^.

### Tissue preparation for histology

Either the entire intestinal tract was removed and divided into small intestine (proximal, middle and distal) and colon, or, in case of wounding/colitis experiments, only the colon was removed. The intestines were flushed with PBS, opened longitudinally using a gut preparation apparatus, and fixed overnight in 10% neutral buffered formalin (Merck). The tissues were then embedded in paraffin and sectioned at 4μm using a microtome (Leica) with DEPC-treated water (Merck) in the water bath. In order to find the ulcers generated by biopsy-wounding, the entire tissue block was sectioned; 4 serial sections collected, followed by a 50μm trim (discarded), throughout the block. H&E staining was performed on one of 4 sections of each series following standard procedures, leaving 3 blank slides per series for other stainings. The mid-ulcer series of sections was determined for all ulcers by identifying their range within the entire set of H&E slides per block. For whole-mount scanning, colons were washed with cold PBS, whole-mounted and fixed in 4% PFA for 3 hours at room temperature.

### *In situ* hybridisation

Standard Advanced Cell Diagnostics RNAscope protocols were followed for both chromogenic ISH (RNAscope^®^ 2.5 HD Assay) and fluorescent ISH (RNAscope^®^ Multiplex Fluorescent Assay v2). For the latter, fluorophores (Cy3 and Cy5) were diluted at 1/1500 in TSA buffer. The following Advanced Cell Diagnostics probes were used: Hs-*BMP4* (454301), Hs-*GREM1* (312831), Hs-*ID1* (414351), Mm-*Clu* (427891-C3), Mm-*Grem1* (314741), Mm-*Id1* (312221), Mm-*Rspo3* (402011), Mm-*Wnt5a* (316791), Mm-*Foxl1* (407401), Mm-*Gli1* (311001), Mm-*Il33* (400590), Mm-*Lgr5* (312171). Chromogenic IHC after ISH was started right after the DAB-washing step in the 2.5 HD assay protocol, fluorescent IHC after ISH was started right after the last HRP-blocking step in the v2 multiplex fluorescent assay protocol. For both staining types, IHC was started with 2 x 5 minutes wash in PBS, followed by 1 hour of blocking. Antibodies were incubated overnight at either room temperature or 4°C.

### Immunohistochemical Analysis

#### Histochemical and immunohistochemical analysis

Sections were de-paraffinized in xylene and rehydrated through graded alcohols to water. Antigen retrieval was done by pressure cooking in 10 mmol/L citrate buffer (pH 6.0) for 5 minutes, for Ki67 followed by 2 minutes incubation in 0.5% Triton-X100 in PBS. Endogenous peroxidase activity was blocked by incubating in 3% hydrogen peroxidase (in water) for 30 minutes (for TdTomato staining hydrogen peroxidase was diluted in methanol instead of water). Next, sections were blocked with 5% serum for 1 hour, after which they were incubated with primary antibodies. Antibodies against the following proteins were used: PTGS2 (BD Bioscience, 610203, 1/50), PDPN (Sino Biology, 50256-RP02-50, 1/500), αSMA (Sigma Aldrich, A2547, 1/1000), pan-cytokeratin (ThermoFisher, 3-9003-82, 1/50), pSmad1/5/8 (Merck, AB3848-I, 1/200), Ki67 (Cell Signaling Technology, CS12202S, 1/500), CHGA (Abcam, ab15160, 1/400), Lysozyme (DAKO, EC3.2.1.17, 1/500), LY6G (BD biosciences, 551459, 1/50), CD45 (Biolegend, 103127, 1/300), F4/80 (ThermoFisher, 14-4801-82, 1/50), CD31 (Cell Signaling Technology, 77699S, 1/100), pHistone3 (Abcam, ab5176, 1/100), TdTomato (Novus Bio, NBP1-96752, 1/500). The sections were then incubated with appropriate secondary antibodies for 30 minutes at room temperature. For chromogenic visualization, sections were incubated with ABC (Vector labs) for 30 min and stained using DAB solution (VectorLabs), after which they were counterstained with hematoxylin, dehydrated and mounted. For immunohistochemical staining after chromogenic ISH, the ImmPRESS-AP polymer reagent and ImmPACT Vector Red AP kit (Vectorlabs) were used to develop a red signal. In case fluorescent secondary antibodies were used, sections were incubated with DAPI (RNAscope^®^ Multiplex Fluorescent kit v2) for 1 minute, then mounted with ProLong Gold antifade mountant (ThermoFisher). For other fluorescent visualizations, Tyramide SuperBoost kits (Alexa Fluor 594 or 647, ThermoFisher) were used. Alcian blue staining was performed standard protocol, briefly, slides were incubated in Alcian Blue for 30 min, and washed in water. They were then incubated in nuclear fast red solution (Sigma) for 5 min and washed in running tap water for 1 min, followed by dehydration and mounting.

### Multiplex Panel

Multiplex (MP) immunofluorescence (IF) staining was carried out on 4um thick formalin fixed paraffin embedded (FFPE) sections using the OPAL™ protocol (AKOYA Biosciences) on the Leica BOND RX^m^ autostainer (Leica, Microsystems). Six consecutive staining cycles were performed using the following primary Antibody-Opal fluorophore pairings; A) Stroma panel: 1) Gremlin 1 (R&D, AF956, 1:750) – Opal 540, 2) CD34 (Abcam, ab81289, 1:3000) – Opal 520, 3) CD146 (Abcam, ab75769, 1:500) – Opal 570, 4) SMA (Abcam, ab5694, 1:1000) – Opal 620, 5) Periostin (Abcam, ab227049, 1:1000) – Opal 690, 6) E-Cad (Cell signalling, 3195, 1:500) – Opal 650. B) Immune panel: 1) Ly6G (BD Pharmingen, 551459, 1:300) – Opal 540, 2) CD4 (Abcam, ab183685, 1:500) – Opal 520, 3) CD8 (Cell signalling, 98941, 1:800) – Opal 570, 4) CD68 (Abcam, ab125212, 1:1200) – Opal 620, 5) FoxP3 (Cell signalling, 126553, 1:400) – Opal 650, 6) E-Cad (Cell signalling, 3195, 1:500) – Opal 690. Primary antibodies were incubated for 60 minutes and detected using the BOND™ Polymer Refine Detection System (DS9800, Leica Biosystems) as per manufacturer’s instructions, substituting the DAB for the Opal fluorophores, with a 10-minute incubation time and without the haematoxylin step. Antigen retrieval at 100ºC for 20 min, as per standard Leica protocol, with Epitope Retrieval (ER) Solution 1 or 2 was performed before each primary antibody was applied. Sections were then incubated for 10 minutes with spectral DAPI (FP1490, Akoya Biosciences) and the slides mounted with VECTASHIELD^®^ Vibrance™ Antifade Mounting Medium (H-1700-10, Vector Laboratories). Whole slide scans and multispectral images (MSI) were obtained on the AKOYA Biosciences Vectra^®^ Polaris™. Batch analysis of the MSIs from each case was performed with the inForm 2.4.8 software provided. Finally, batched analysed MSIs were fused in HALO (Indica Labs), to produce a spectrally unmixed reconstructed whole tissue image, ready for analysis.

### Whole-mount antibody staining

Colon was sectioned into 2-3 cm pieces, washed in 0.1% PBS-Tween for 2 days, and blocked in 10% donkey serum in PBS overnight at 4°C, incubated with DAPI (10 μg/mL stock) in 10% donkey serum in PBS for 3 days. The tissue was finally washed with PBS-T for 1 day before being imaged on a TCS SP5 confocal microscope (Leica).

### Organoid Culture

Organoid cultures were started and maintained principally as described by Sato et al (Sato et al., 2009), except that Noggin was replaced by 0.1μg/mL recombinant GREM1 protein (R&D Systems). The culture medium to maintain the organoids was Advanced DMEM/F12 (Thermofisher) supplemented with Penicillin/Streptomycin, 10mM HEPES, 1x Glutamax, 1x B27 supplement (Thermofisher), 1.25mM N-Acetyl-L-cysteine (Sigma), (50 ng/mL) rmEGF (Thermofisher), RSPO1, WNT3A (termed ERW-medium) and GREM1. The RSPO1 source was 10% conditioned medium. The WNT3A source was 50% conditioned medium from L cells (ATCC CRL-2647) prepared following the company’s protocol. Small intestinal organoids isolated from steady state WT mice were mechanically chopped into small pieces using a small-orifice Pasteur’s pipette and seeded in either Matrigel (Corning) or collagen matrix. Collagen matrix was prepared on ice by adding 10x PBS (to make 1x PBS) and 1N NaOH (to make 25mM NaOH) to Collagen I (Rat tail, Thermo Fisher). Organoids were treated with ERW medium, ERW medium with GREM1, or ERW medium with different concentrations of recombinant mouse Bmp4 protein (R&D systems). The organoids received new medium after 2 days and were imaged and harvested for RNA analysis after 4 days of treatment.

### Quantification and statistical analysis

#### Imaging and staining quantifications

Fluorescent pictures were taken with a Nikon TE2000u equipped with a Hamamatsu C4742-96 camera. Organoid pictures were taken with a stereomicroscope (Leica S8 AP0) equipped with a camera (Chromyx). Pictures of chromogenically stained sections were taken using the Aperio CS2 slide scanner. HALO^®^ software (Indica Labs) was used for quantification of *Grem1*-staining at ulcers. Regions were drawn demarcating the ulcer bed and the first 0.5mm of colon tissue on either side of the ulcer, the latter further separated into muscularis propria, muscularis mucosa and lamina propria. *Grem1* staining positivity was measured as percentage area of each region. Quantifications of other stainings were done using Qupath open source digital software. Annotation objects demarcating ulcer-adjacent crypts and WAE were drawn and cells were automatically detected by Qupath. The nuclear or cellular mean DAB optical densities (mDOPs) of all cells were normalized by subtracting the lowest detected nuclear or cellular mDOP in that image. Cells with normalized mDOPs above a set threshold per staining target were counted as positive. For Alcian Blue staining quantification, "Haematoxylin" values were set to red color values and “Stain 2” to blue color values to adapt to the different chromogens used. The counting of positive crypts in *en face* sections of DSS-treated animals was done manually, aided by automatic positivity labelling of individual cells by Qupath. For *Bmp4*-ISH quantification, extra care was taken to discriminate between epithelial and stromal signal source and any epithelial cells with two or more subcellular DAB spots were regarded positive, while for tdTom-IHC quantification, any cells with nmDOP >0.1 were regarded positive. Crypts with more than half of its cells being positive were counted as positive crypts. The counting of fully clonal crypts at steady state conditions was done manually. The fraction of traced cells per cell position in the crypt as well as the number of traced cells per villus were manually quantified independently by 3 different observers, and their values were averaged. For quantifying fluorescence images, the mean cellular pixel values of all detected cells were normalized per channel by subtracting the lowest detected mean cellular pixel value. Cells with normalized mean cellular pixel value above a set threshold per channel were counted as positive. Quantification of co-stainings of tdTomato-IHC or *Bmp4*-ISH with Ki67 or differentiation markers was done manually in Qupath. Detection of overlap with the *Grem1*-ISH staining pattern (Figure S2) was done using the “Colocalization Threshold” option in ImageJ. Quantification of organoid area was performed using the plugins “Adaptive Thr” and “MorphoLibJ” in ImageJ.

#### Colon ulcer measurements

Ulcer width was measured as the distance between the two ulcer-adjacent crypts, following the curve of the muscularis mucosa, if present. WAE-length was measured as the length of the epithelium covering the ulcer bed until its transition into the ulcer-adjoining crypt. The degree of ulceration was determined by measuring the total length of denuded regions over the total distal colon length, from the proximal folds down.

## AUTHOR CONTRIBUTIONS

MAJK, HD and SJL conceived and designed the project. Funding obtained by SJL. Experiments were conducted by MAJK, HD, NN, EM, MC, SB, MF, LL. Data provision and bioinformatic analysis carried out by GV, AA. Pathology support, image analysis, tissue provision and intellectual input from VHK, LMW, JEE. Conceptual input and data interpretation AS, DW. Manuscript written by MAJK, GV and SJL.

## DISCLAIMER

The views expressed are those of the author/s and not necessarily those of the NHS, the NIHR or the Department of Health

