## Supplementary Figures and Tables for "BMP pathway antagonism by *Grem1* regulates epithelial cell fate in intestinal regeneration"

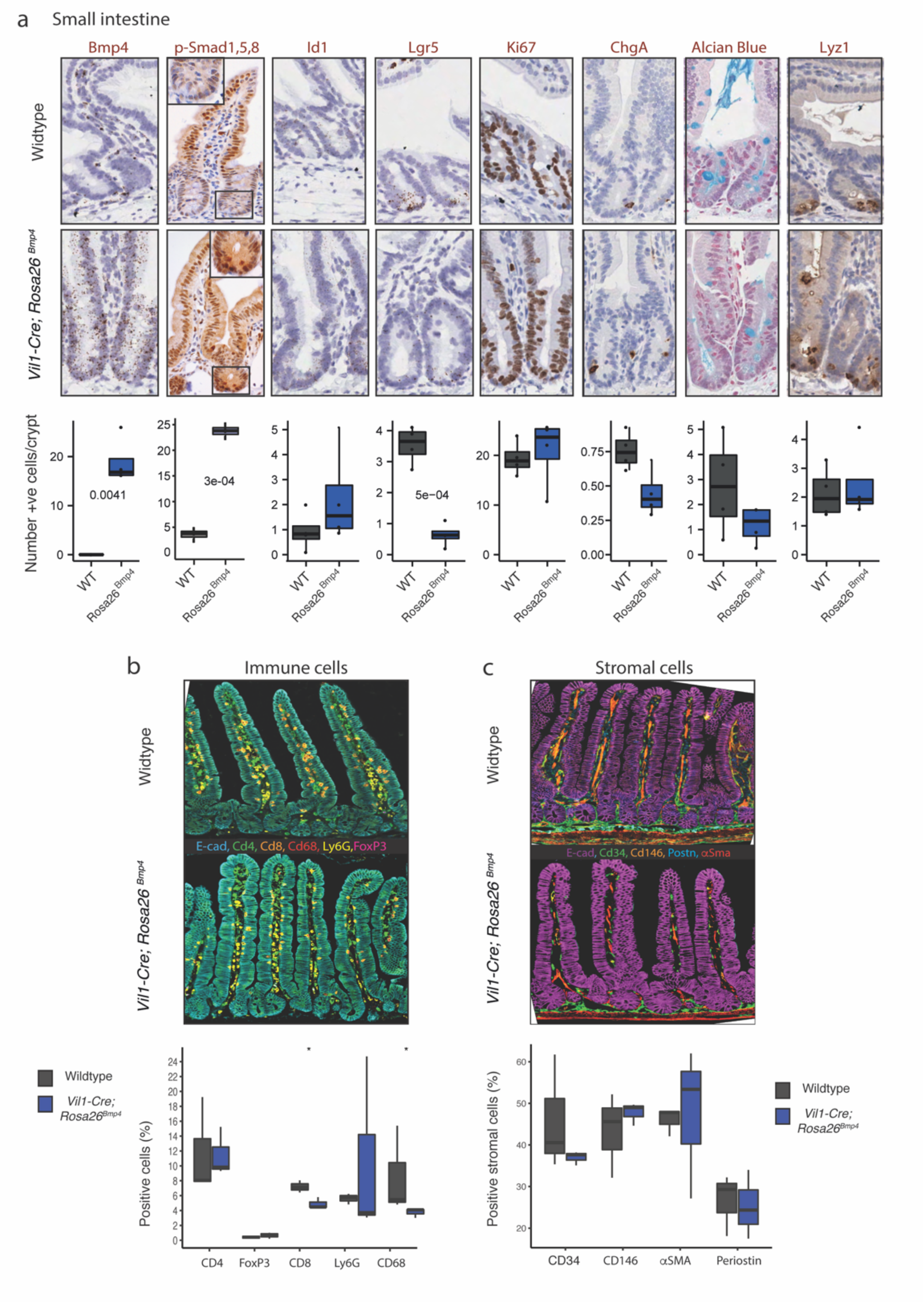


**Supplementary figure 1. Steady-state *Vil1-Cre;Rosa26$^{Bmp4}$* mouse small intestinal phenotype a.** ISH/IHC phenotyping and cell quantification of small intestine in steady state *Vil1-Cre;Rosa26$^{Bmp4}$* and control mice. (t-test, n=4 mice per genotype). **b,c.** Multiplex IHC and cell quantification to show small intestinal **b.** immune and **c.** stromal cell landscapes in wildtype and *Vil1-Cre;Rosa26$^{Bmp4}$* animals (* p<0.05, t test, n=3 per genotype)

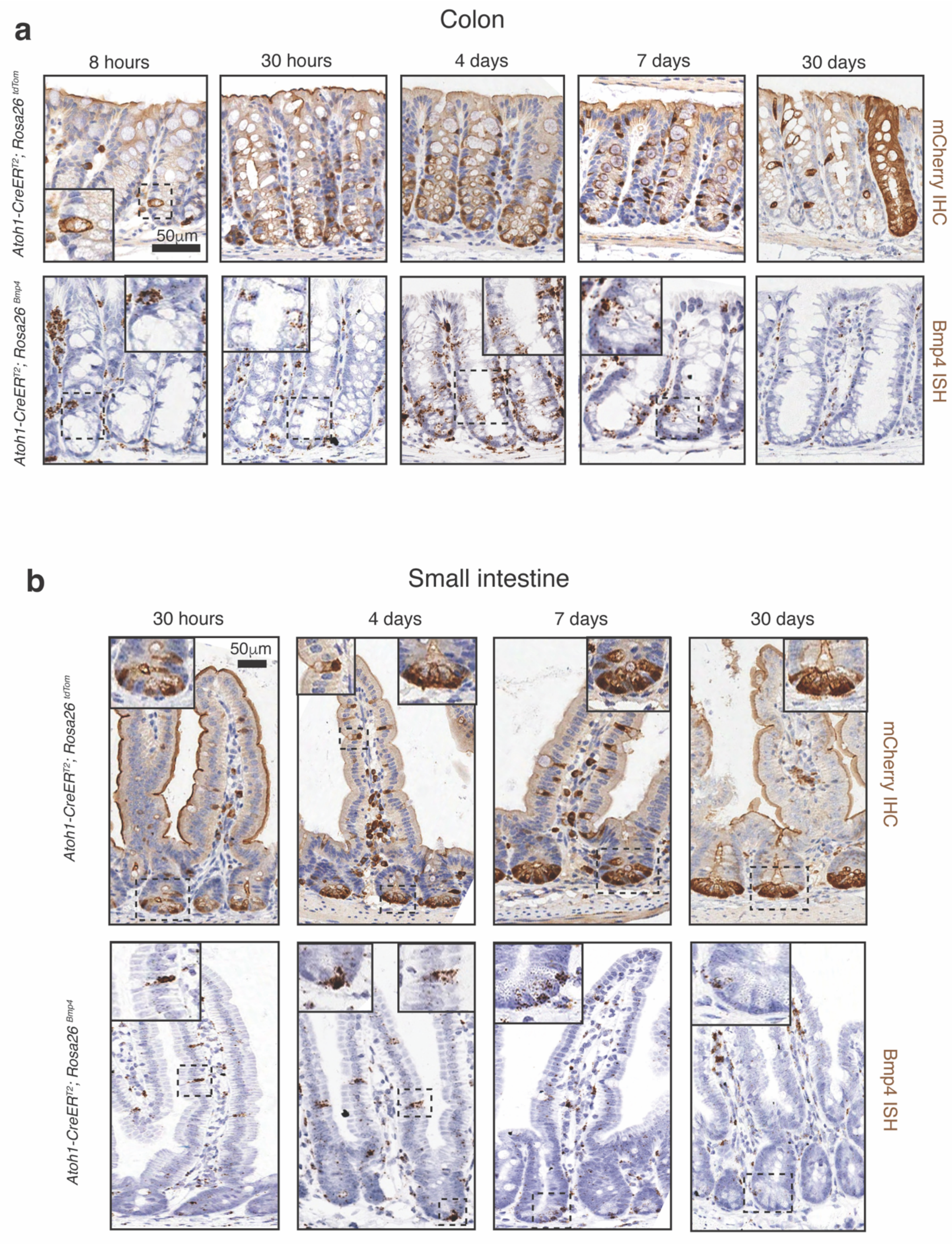


**Supplementary figure 2. Staining for cell counting.** Representative images of tdTomato IHC (brown) and *Bmp4* ISH (brown dots) in secretory epithelial cells of the **a.** colon, and **b.** small intestine at different time points after recombination in *Atoh1CreER*$^{T2}$*;Rosa26*$^{tdTomato}$ *and Atoh1CreER*$^{T2}$*;Rosa26*$^{Bmp4}$ mice. Scale bar 50μm all panels.

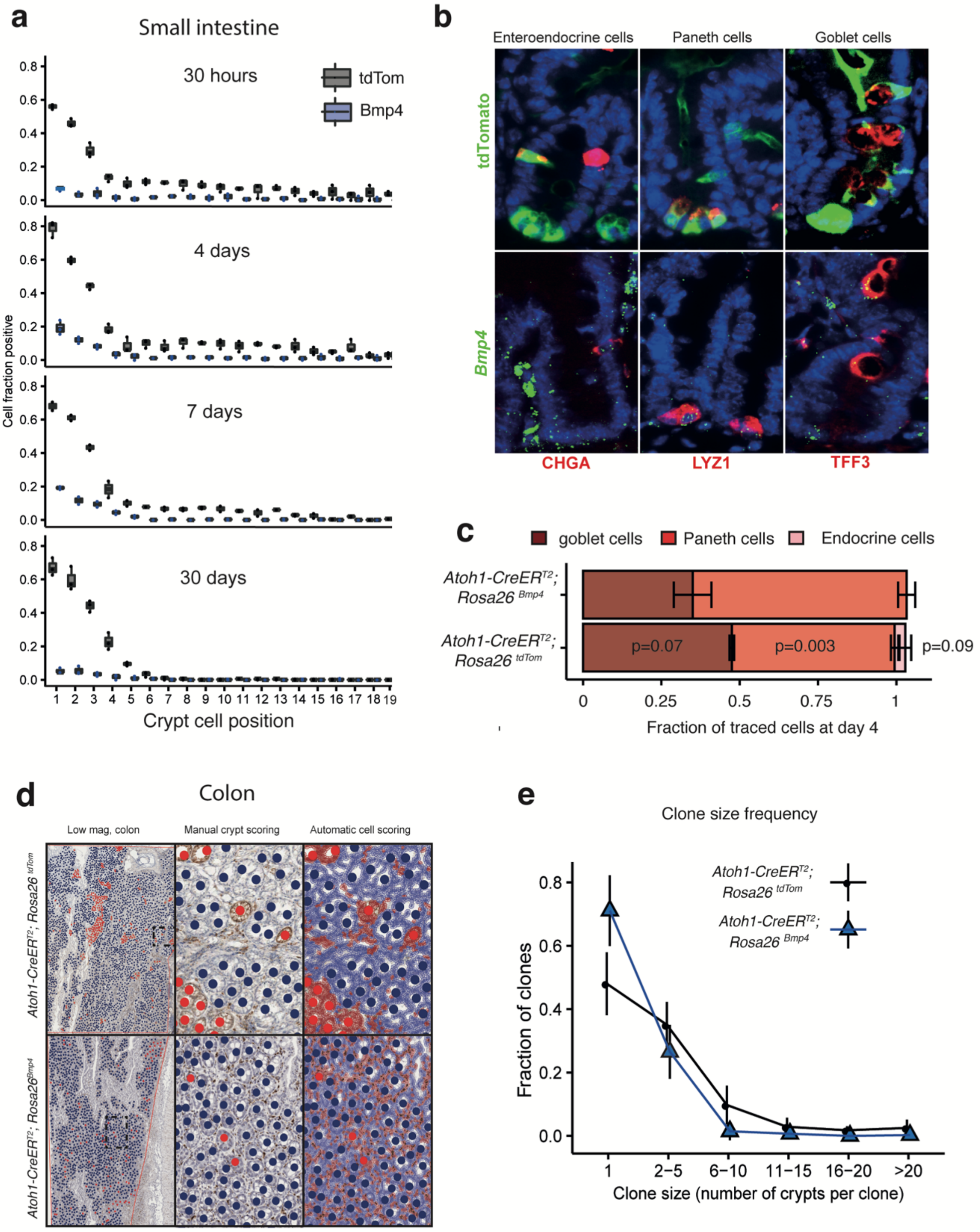


**Supplementary figure 3. Secretory cell model assessment. a.** Steady state small intestinal crypt cell position counts of tdTomato IHC (grey) or *Bmp4* ISH (blue) stained cells over time following recombination (n=3 mice, 7 days: n=2 mice, 30 crypts counted per mouse) **b.** Co-stain of tdTomato IHC (green, top panels) or *Bmp4* ISH (green, bottom panels) with IHC markers for enteroendocrine cells (Chromogranin A, red), Paneth cells (Lysozyme, red) and goblet cells (TFF3, red) in the small intestine secretory cell models in steady state, 4 days after recombination. **c.** Quantification of fraction of traced cells co-staining for individual secretory cell markers in different secretory cell models in steady state, 4 days after recombination (t-test, between the same cell types in each genotype, p values as stated, n=3 mice, 9-22 crypts counted per mouse) **d.** Representative images of *en face* sections from secretory cell models taken 30 days after initiation of DSS colitis. Low power images show variable expansion of clonal patches in different genotypes. High power images demonstrate manual and automated digital pathology (QuPath) scoring of positive epithelial cell staining, used to exclude non-contributory stromal cell staining. **e.** Quantification of clone size frequency in different secretory cell models, 30 days after DSS initiation shows greater fraction of multi-crypt and large clonal patches in $Atoh1CreER^{T2};Rosa26^{tdTom}$ animals (Rosa26Bmp4 n=5 mice, Rosa26tdTom n=4 mice). ** $P < 0.01$.

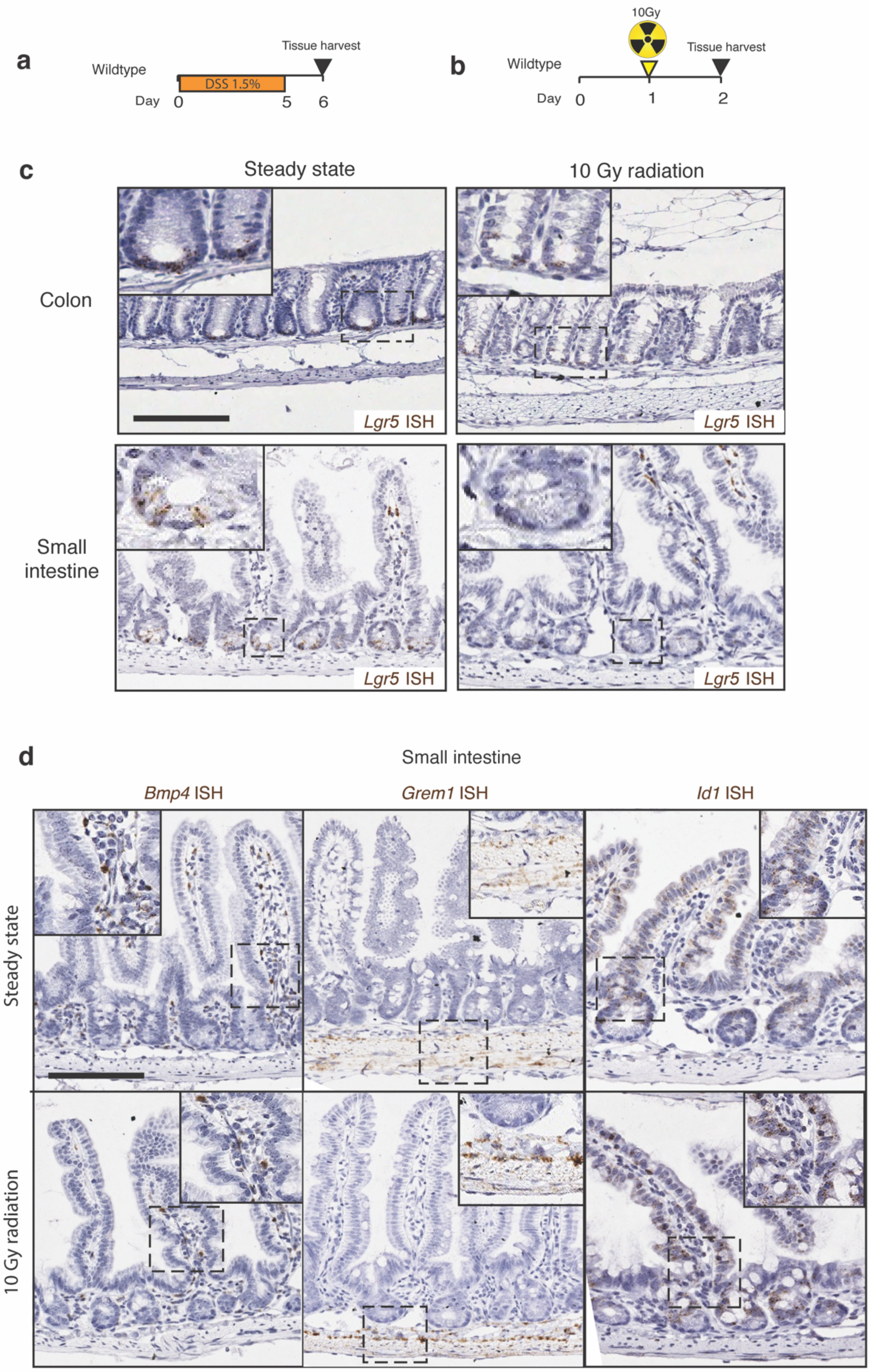


**Supplementary figure 4. Impact of 10Gy intestinal regeneration.** Schematic showing schedule of **a.** DSS administration and **b.** intestinal irradiation and tissue harvesting in BMP signalling disruption animal models**. c.** *Lgr5* ISH to show loss of *Lgr5* expression in crypt base columnar stem cells 24 hours after 10Gy whole body irradiation **d.** *In situ* hybridization of *Bmp4, Grem1* and *Id1* in mouse small intestine in steady state and following 10Gy irradiation. Scale bars: 200*µ*m, magnification applies to all images, except insets.

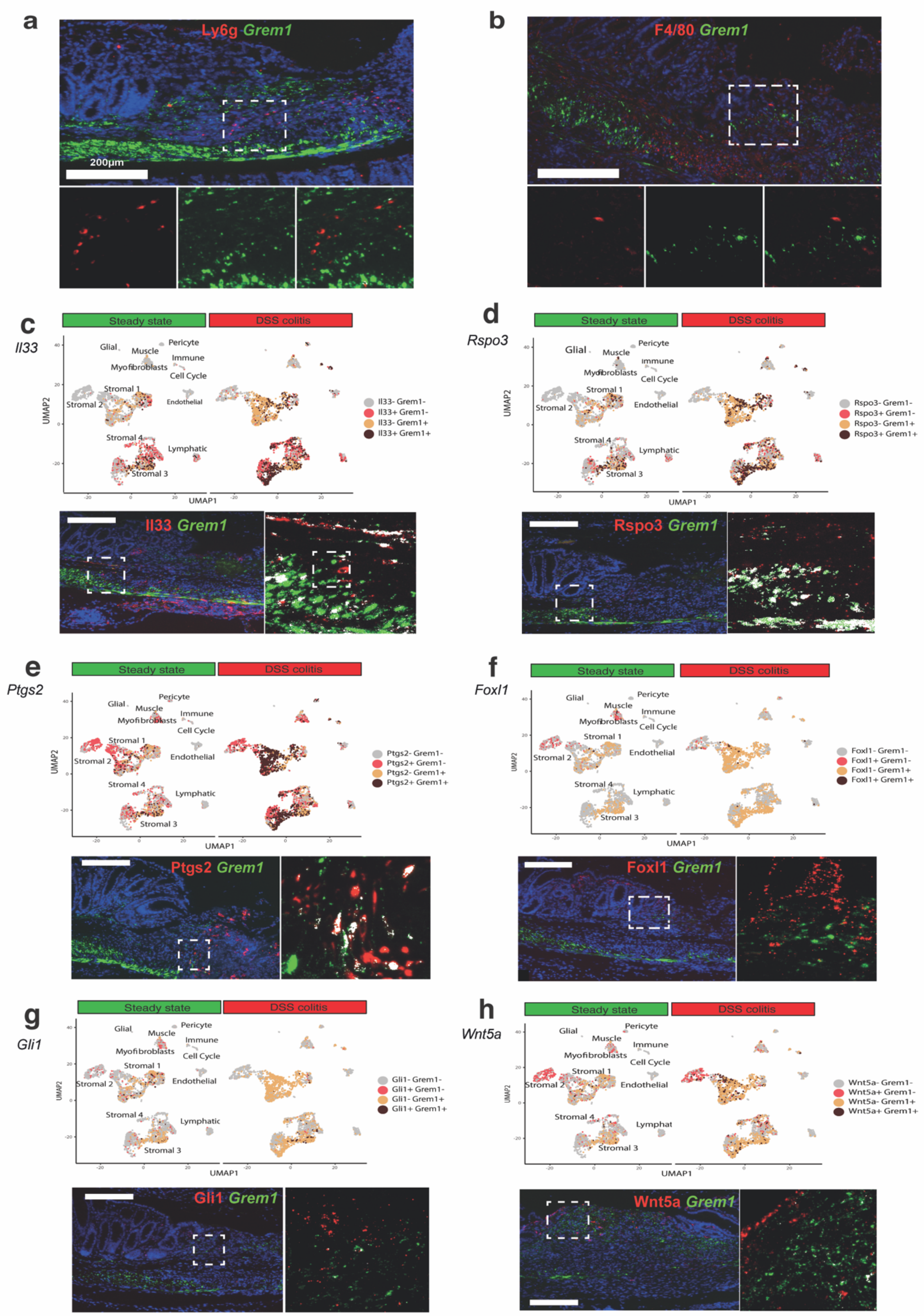


**Supplementary figure 5. Co-expression of *Grem1* with intestinal mucosal immune and stromal cells a,b.** There was no overlap of fluorescent expression of *Grem1* mRNA (green)

with **a.** neutrophils marked by LY6G (red), or **b.** with macrophages marked by F4/80 (red). **c-e.** Single cell transcription U-MAP plots (steady state and DSS colitis) and co-ISH/IHC expression for *Grem1* (green) with **c.** *Il-33* (red) **d.** *R-Spo3* (red) and **e.** Ptgs2 (anti-COX2 antibody - red) showing overlap of expression (dark brown dots on U-MAP, white colour on IHC/ISH) in discrete cells of the muscularis or ulcer bed in intestinal ulcers. **f-h.** Single cell transcription U-MAP plots (steady state and DSS colitis) and co-ISH showing predominantly distinct cell populations expressing *Grem1* (green) and **f.** *Foxl1* (red) **g.** *Gli1* (red) and **h.** *Wnt5a* in intestinal ulcers. Scale bar 200*μ*m in all panels.

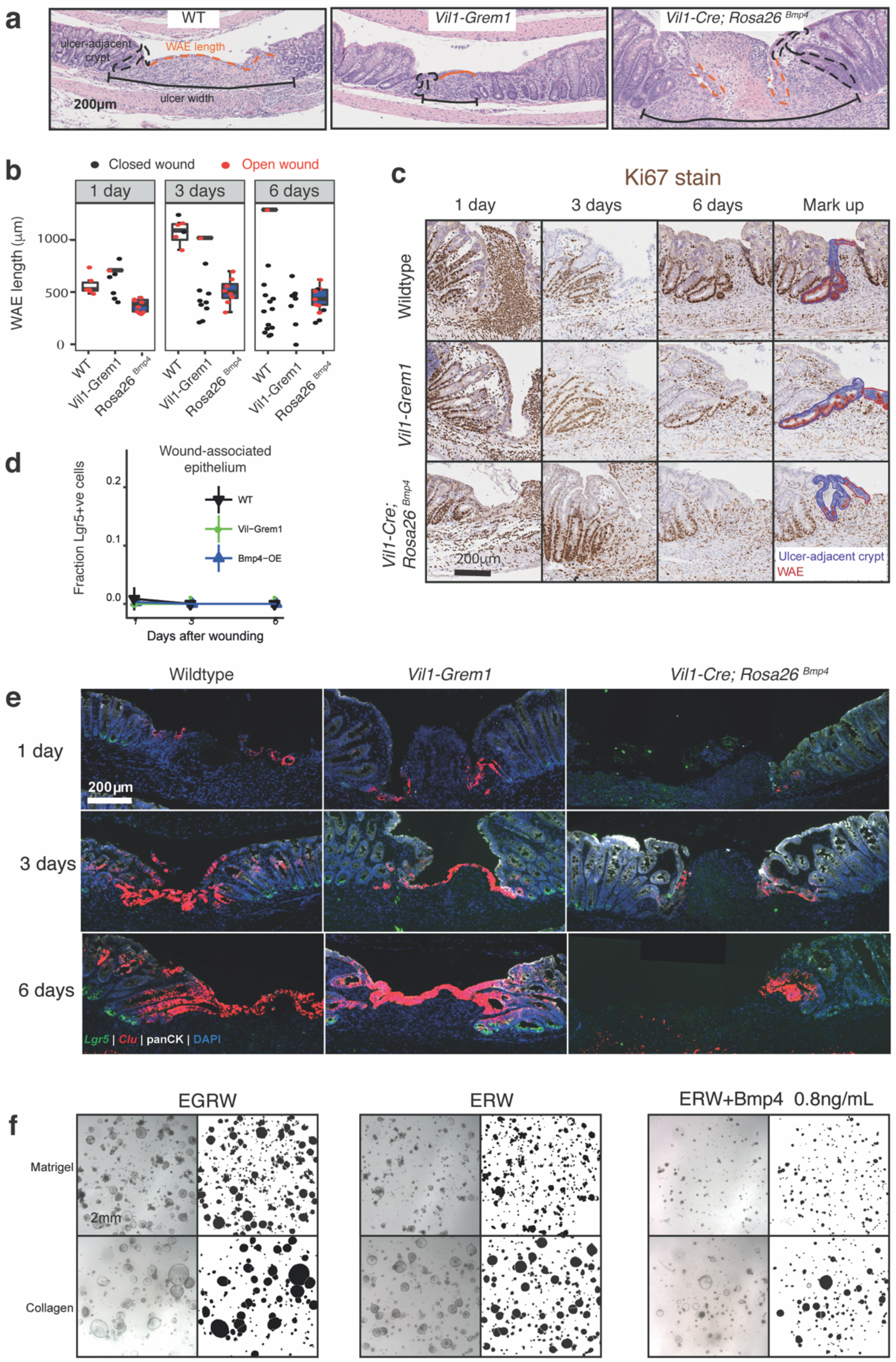


**Supplementary figure 6. Ulcer quantification and stem cell phenotyping. a.** Representative H&E images of 3 day old biopsy wounding ulcers in different genotype animals, containing examples of ulcer-adjacent crypts (dashed black lines), ulcer width (solid black lines), and wound associated epithelium (WAE) length (dashed orange lines). **b.** Dot and box plot of WAE length over time in endoscopy biopsy ulcers from WT, *Vil1-Grem1* and *Vil1-Cre;Rosa26*$^{Bmp4}$ animals. Ulcers completely covered with WAE ("Closed", black dots) were not included in the boxes, while incompletely covered ulcers ("Open", red dots) are shown as both dots and boxes (n=6-15 ulcers). **c.** Cell staining quantification reveals absence of *Lgr5* expression in wound associated epithelium over time in all mouse genotypes (n=3-6 mice per group). **d.** Representative images of Ki67 stain in different mouse genotypes, with digital pathology cell identification mark up (QuPath) used for quantification of cell proliferation in ulcer-adjacent crypts. **e.** Representative images of ISH for *Lgr5* (green), Clusterin (red) with co-stained IHC for pan-cytokeratin (white) and DAPI (blue) in the endoscopic biopsy wounds of different genotype animals over time. **f.** Stereomicroscopy images with organoid area quantification of organoids grown in matrigel or collagen and treated with media containing variable recombinant proteins (E: Epidermal growth factor, G: Gremlin1, R: R-spondin1, W: Wnt3a). Scale bars 200$\mu$m all panels.

**Supplementary Table S1 -** Taqman probes used in qRT-PCR experiments.

| Gene | TaqMan probe | Source | Identifier |
|---|---|---|---|
| Bmp2<br>Bmp4<br>Bmp5<br>Bmp7<br>Chrdl1<br>Chrdl2<br>Clu<br>Gli1<br>Grem1<br>Grem2<br>Id1<br>Il6<br>Lgr5<br>Nog<br>Serpine | Mm01340178_m1<br>Mm00432087_m1<br>Mm00432091_m1<br>Mm00432102_ml<br>Mm00473158_m1<br>Mm01136674_m1<br>Mm01197002_m1<br>Mm00494654_ml<br>Mm00488615_s1<br>Mm00501909_m1<br>Mm00775963_g1<br>Mm00446190_m1<br>Mm00438890_m1<br>Mm01297833-s1<br>Mm00435858_ml | Life Technologies | Cat# 4331182 |
| Gapdh | | Life Technologies | Cat# 4352661 |
| BMP2<br>BMP4<br>BMP5<br>BMP7<br>CHRDL1<br>CHRDL2<br>GREM1<br>GREM2<br>ID1<br>LGR5<br>NOG | Hs00154192_m1<br>Hs00370078_m1<br>Hs00234930_m1<br>Hs00233476_m1<br>Hs00292767_m1<br>Hs00248808_m1<br>Hs00171951_m1<br>Hs03986140_s1<br>Hs03676575_s1<br>Hs00173664_m1<br>Hs00271352_s1 | Life Technologies | Cat# 4331182 |
| GAPDH | | Life Technologies | Cat# 4333764F |
